# A rational approach for the targeted discovery and characterisation of microbiome-derived therapeutics

**DOI:** 10.1101/2024.10.22.619608

**Authors:** Lutz Krause, Joel Boyd, Charlotte Vivian, Annika Krueger, Jeimy Jimenez Loayza, Johanna K. Ljungberg, Joyce Zhou, Andrea Rabellino, Bozica Nyeverecz, Michael Nissen, Kaylyn Tousignant, Mareike Bongers, Martha M. Cooper, Donovan H. Parks, Areej Alsheikh, Patricia Vera-Wolf, Mitchell Sullivan, Rhys Newell, Kelly-Anne Masterman, Peter Evans, Liang Fang, Samantha MacDonald, Maria Chuvochina, Darrell Bessette, Kate Zimmermann, Alena Pribyl, Hiram Chipperfield, Shandelle Caban, Huw McCarthy, Joanne Soh, Luke Reid, Ian H. Frazer, Nicola Angel, Tony Kenna, David L.A. Wood, Blake Wills, Jakob Begun, Simon Keely, Trent Munro, Gene W. Tyson, Philip Hugenholtz, Páraic Ó Cuív

**Author notes:** Corresponding authors Lutz Krause, Páraic Ó Cuív.

## Abstract

The human gut microbiome is intrinsically involved in health and disease, representing a wealth of untapped therapeutic potential. Here, we demonstrate the utility and potential of a metagenome guided, large cohort-based approach for the rational selection of live biotherapeutics from the human gut. We applied this approach to Inflammatory Bowel Disease (IBD), identifying several lead candidates that were significantly depleted in individuals with IBD compared to healthy controls. Their therapeutic potential was assessed in preclinical models of IBD where they improved markers of disease pathology by reducing inflammation and promoting mucosal healing and wound repair. All leads had excellent safety profiles *in silico* and *in vitro*, and several additionally presented favourable manufacturing properties, supporting their progression into clinical trials. We believe that this rational approach will be generalisable to any disease state with underlying microbiome aetiology and will expedite the development of novel microbiome-derived therapeutics to improve human health.

## Introduction

The human gut microbiome influences numerous host functions, ranging from metabolism and immune development to cognitive function and behaviour^1–3^. The intestinal microbiome has been implicated in common chronic diseases including inflammatory disorders^4^, metabolic diseases^5^, neurodegenerative diseases^6^, and cancer^7^, offering further evidence that the microbiome is critical for maintaining host homeostasis. Consequently, the development of new therapeutics that aim to restore health-promoting microbial functions holds great potential.

There have been two recent Food and Drug Administration (FDA) approvals for human faecal donor derived, complex consortium-based microbiome products for treatment of recurrent *Clostridioides* (formerly *Clostridium*) *difficile* infection^1,2,8,9^. However, despite extensive study of the gut microbiome over the past decade, development of non-donor derived, rationally designed effective microbial therapeutics has remained elusive. Previous attempts to identify therapeutics from the gut microbiome have relied on 16S rRNA gene sequencing, low resolution metagenomic sequencing, or isolation and screening approaches, where culture biobanks are screened for activities of interest, such as transcriptional regulation^10,11^, anti-inflammatory^12,13^ or neuroprotective effects^14^. Without disease context from high-resolution human datasets, these methods have limited probability of identifying the most relevant microorganisms capable of treating or preventing disease. This is confounded by the fact that an estimated 63% of microbial species from the human gut have yet to be cultured in the laboratory^15^. Existing biobanks therefore only provide access to a fraction of the potential therapeutic pool.

Here we present a metagenome-guided, large cohort-based approach for the rational selection of live biotherapeutic (LBP) leads from amongst the thousands of microbial species known to inhabit the human gut. We applied the approach to Inflammatory Bowel Disease (IBD), a chronic inflammatory disorder of the gastrointestinal tract affecting ∼6.8 million people annually^16^, with focus on the two major disease subtypes, Crohn’s disease (CD) and ulcerative colitis (UC). Existing treatments aim to control the aberrant immune response characteristic of IBD; however, response rates remain relatively low, and with current standard of care, the cumulative risk of relapse is 90% in patients with UC and CD after 10–25 years of follow up^17,18^. Furthermore, there is a clinical need for new therapeutics that induce mucosal healing as this is associated with improved disease outcomes^19,20^, and is a key unmet therapeutic need. Using our approach, we discovered multiple promising therapeutic leads enriched in healthy individuals relative to IBD patients, including several previously uncultured species that were obtained by genome-directed isolation (GDI)^21^. Our leads suppress disease-associated inflammatory pathways and reproducibly promote mucosal healing and wound repair in mouse disease models and human gut epithelial cell-based assays. These LBP candidates may provide the basis for long-term, safe treatment of IBD. Our results demonstrate that a rational data-guided approach is a reliable means for the expedited discovery of LBPs that should be generalisable to many other disease states.

## Results

We developed a drug discovery and development platform to obtain novel LBP candidates, using IBD as an exemplar disease (Figure 1). The platform uses the Microba Community Profiler (MCP) and associated reference genome database^22,23^ that together allow the accurate taxonomic and functional characterisation of gut microbiomes. After accounting for confounding factors by matching disease and control groups, candidate therapeutic leads that show the strongest associations with health and disease are selected using machine learning and statistical techniques. Healthy donor faecal samples with a naturally high abundance of target species are then used to isolate leads in pure culture. Previously uncultured species are obtained by GDI^21^, which involves metabolic modelling to design media that selectively promote the growth of target species. Following optimisation of growth conditions, target species are characterised through a series of *in vitro* and *in vivo* disease assays to evaluate their potential for therapeutic development (Figure 1).

**Figure 1.**
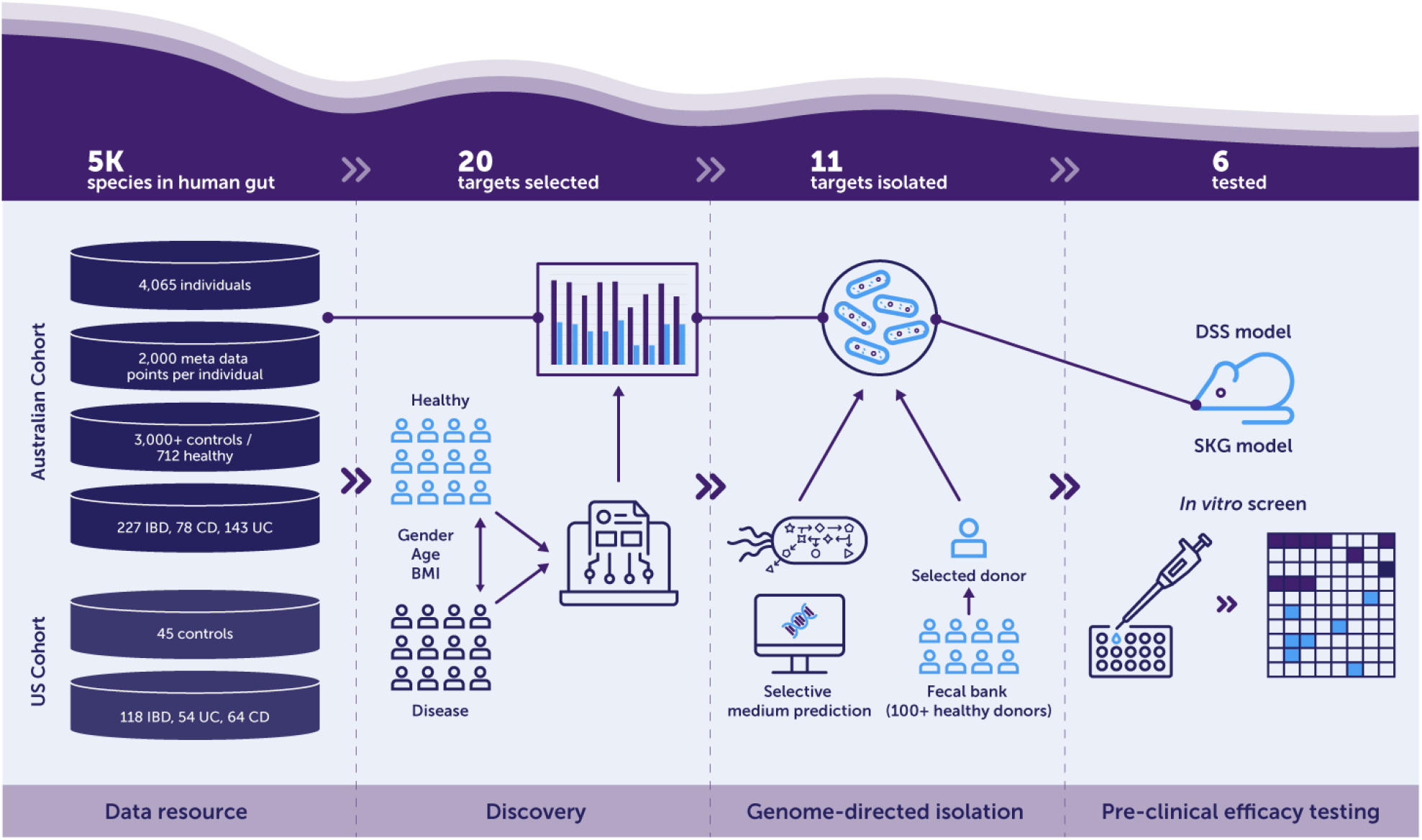
Overview of a rational approach for the discovery and characterisation of novel microbiome-derived therapeutics. Microbial lead species are identified through comparative analysis of faecal metagenomes from large human cohorts. Leads are then isolated using genome-directed isolation, and their therapeutic potential assessed in animal disease models. Disease relevant functional activities and potential mechanisms of action are further characterised using *in vitro* assays. Here, the approach was applied to the discovery of microbial therapeutics for the treatment of IBD.

### Data driven discovery of targets for IBD therapeutics

IBD leads were identified by comparing the gut microbiomes of healthy individuals to IBD cases. The primary data used for this analysis were stool metagenomes from 241 healthy and 80 IBD participants (30 CD; 50 UC), selected from 4,065 metagenomes obtained from a cross section of the Australian population (Figure 1). Healthy controls and disease study groups were matched by age and gender, resulting in cohorts of 231 healthy and 79 IBD, 150 healthy and 30 CD, and 147 healthy and 49 UC (Figure S1). Consistent with prior research^24–26^, the gut microbiomes of both major subtypes of IBD were characterised by significantly reduced species diversity and richness compared to controls, and a strong microbial signature discriminating disease and control groups (Figure 2A). We also replicated previously reported findings for several disease-associated species, including an increased prevalence of *Enterocloster* (formerly *Clostridium*) *bolteae* and *Hominicoccus* (*gen. nov*., formerly *Ruminococcus*) *gnavus*, and decreased prevalence of *Faecalibacterium prausnitztii* and *Roseburia hominis* in IBD relative to controls (data not shown)^24–26^. We next conducted a meta-analysis combining our Australian cohort with a US-based IBD cohort^25^ (45 controls, 118 individuals with IBD, including 64 CD, and 54 UC), which we re-analysed using the MCP profiler. Our analyses identified 20 bacterial lead species that were consistently detected in the healthy cohort but significantly depleted in individuals with IBD (Figure 2B). Many of the lead species had not previously been associated with IBD and we attribute the identification of these additional leads to the precision of the MCP^22^ and comprehensiveness of its reference genome database.

**Figure 2.**
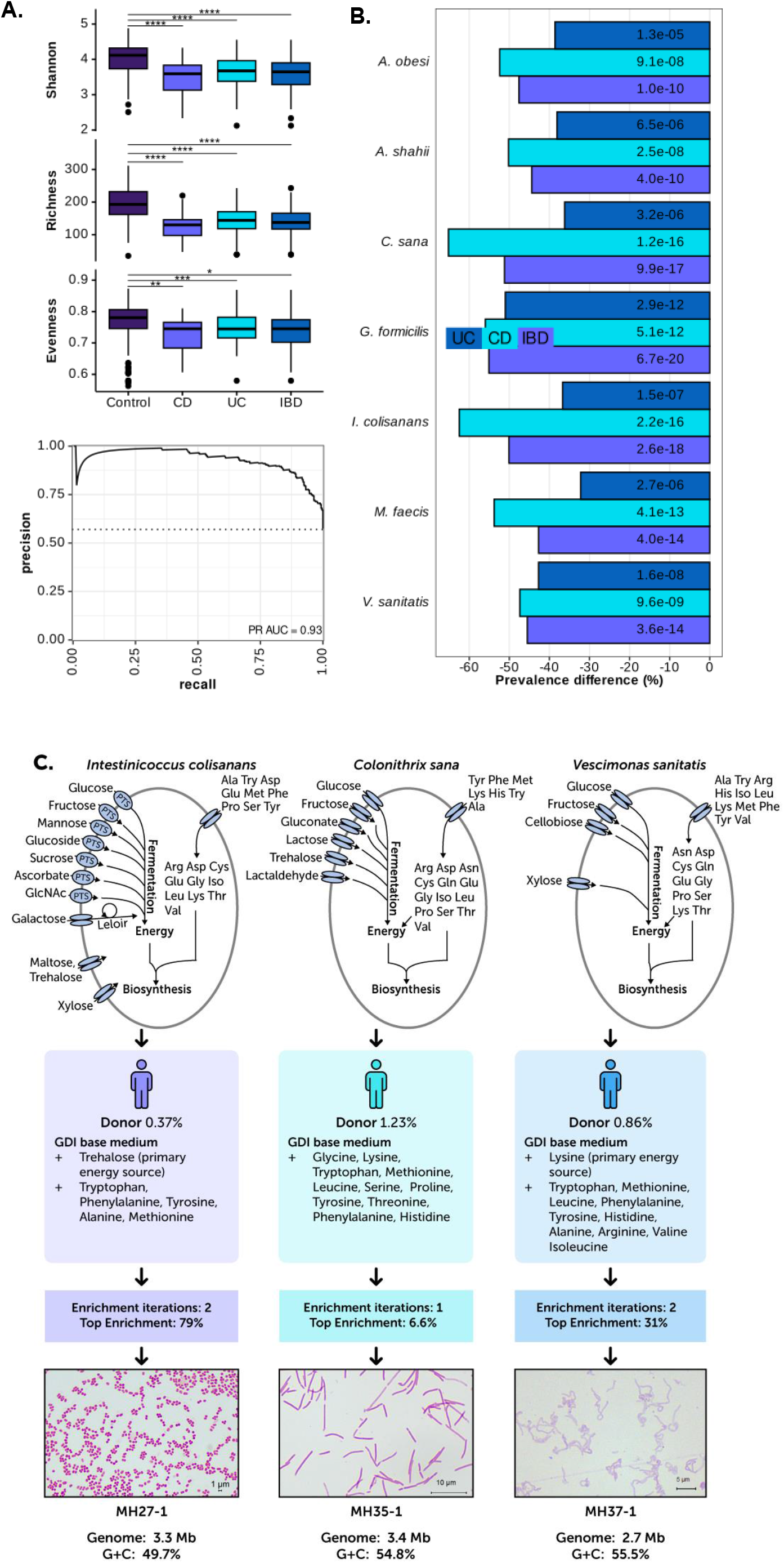
(A) Microbial alpha diversity for human healthy adults (n=231), subjects diagnosed with IBD (n=79), CD (n=30), and UC (n=49) in a large Australian cohort (t-test versus controls; **** = FDR <0.0001; *** = FDR <0.001; ** = FDR <0.01; * = FDR <0.05). Precision-recall curve for a gradient boosting machine learning model discriminating controls from individuals with IBD in a combined Australian and US cohort. B) Percent reduction in prevalence of identified lead species in IBD, UC and CD compared to controls across a combined cohort from Australia and the US. FDR corrected Cochran-Mantel-Haenszel P-values are shown in bars. (C) Genome-directed isolation (GDI) of three previously uncultured species. Shown are predicted carbon and amino acid sources that were added to the base medium for each target species. We also show the prevalence of each strain in the healthy donor sample used to isolate the strain, the number of enrichment iterations, and the percent enrichment after growth on GDI designed media. Following enrichment, pure isolates were obtained through dilution to extinction.

### Isolation of therapeutic leads

We next isolated strains from the identified lead species to assess their therapeutic activities. Notably, 14 of our 20 lead species had not been previously cultured to our knowledge at the time, highlighting the untapped therapeutic potential of the gut microbiome. To isolate these uncultured species, we designed media that selectively promote growth of target organisms using GDI, and obtained axenic cultures of three previously uncultured species; UBA1417 sp003531055 (strain MH27-1), ER4 sp000765235 (strains MH35-1 and MH35-2) and *Vescimonas* sp000435555 (strain MH37-1) (Figure 2C). Species names are listed according to the Genome Taxonomy Database (GTDB) release 08-RS214^27^. Since obtaining these isolates, another strain each of UBA1417 sp003531055 and ER4 sp000765235 were isolated and validly named *Hominenteromicrobium mulieris* (for which we proposed the synonym *Candidatus* Intestinicoccus colisanans^28^) and *Hominiprocola fusiformis*, respectively^29^. Customised broth media were inoculated with healthy donor faecal samples containing the target species, which were then serially diluted to extinction, and axenic liquid cultures were obtained after only one or two enrichment iterations (Figure 2C; see Methods and Supplementary Materials). *H. mulieris* strain MH27-1 was isolated based on its predicted ability to use trehalose and inability to synthesise several amino acids including all three aromatic amino acids. ER4 sp000765235 strains MH35-1 and MH35-2 were isolated based on their predicted use of selected amino acids as an energy source. *Vescimonas* sp000435555 strain MH37-1, for which we propose the name *Vescimonas sanitatis* sp. nov (see taxonomic description below), was obtained based on its predicted use of lysine as an energy source and a number of amino acid auxotrophies. According to the GTDB^23^, *V. sanitatis* is closely related to the recently described human faecal isolates *Vescimonas coprocola* and *Vescimonas fastidiosa*^30^. More details on the isolation of these strains are provided in the Methods and Supplementary Materials. Taxonomic descriptions are provided at the end of this manuscript. Separately, we isolated new strains of *Alistipes shahii* (MH21-1), *Alistipes communis* (MH22-1), *Mediterraneibacter faecis* (MH23-1, MH23-3) and *Gemmiger formicilis* (MH32-1) using established culturing approaches (e.g.^31^, see Methods and Supplementary Materials). The nine isolated leads span seven species, six genera, five families and two phyla (Figure 3). With the exception of *V. sanitatis* MH37-1, all isolates grew well under laboratory conditions thereby enabling an assessment of their therapeutic potential.

**Figure 3.**
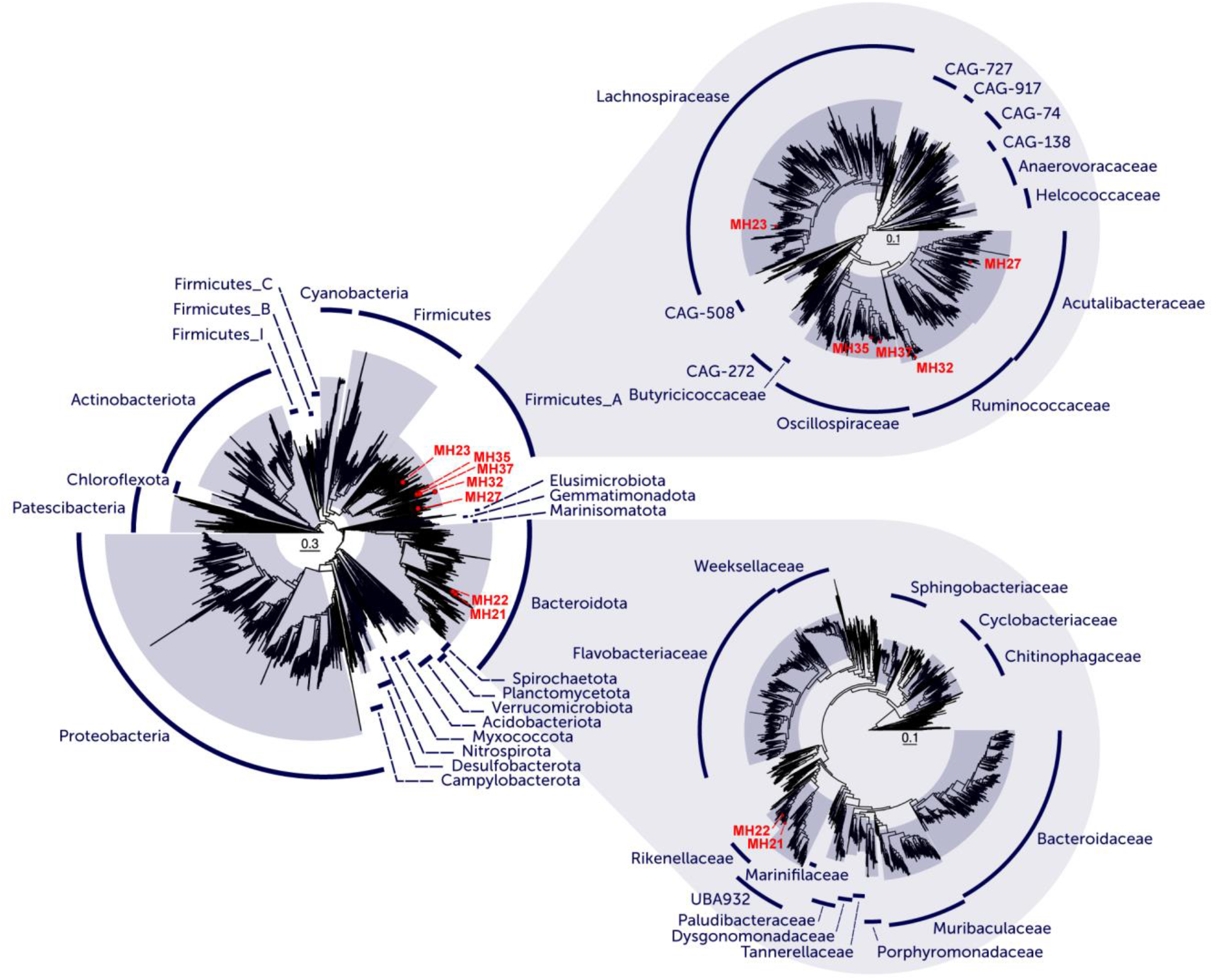
Phylogenetic tree highlighting the phylogenetic diversity of identified IBD leads (based on GTDB r89).

### Therapeutic leads ameliorate murine colitis and ileitis

The prophylactic and therapeutic potential of all bacterial leads except *V. sanitatis* MH37-1 was evaluated *in vivo* using animal models of IBD. The dextran sulphate sodium (DSS) induced murine colitis model, a well-documented model of epithelial injury and repair^32^, was used in a blinded prophylactic study (Figure 4A). Live cultures of *A. shahii* MH21-1, *M. faecis* MH23-1, *H. mulieris* MH27-1, and *G. formicilis* MH32-1, and an equal parts per volume combination of these four leads (MAP#4 consortium) were administered by oral gavage once daily for eight days at 2×10^8^ cells per mouse, starting one day prior to DSS treatment. The steroid prednisone was used as a positive control as it is a mainstay of IBD treatment during flares and effective in the DSS model^33^. DSS treatment resulted in a significant reduction in body weight (Figure S2B), which is an accurate and reliable indicator of colitis^34^. Prednisone exacerbated the DSS-induced weight loss in this model, as expected^35^, whereas treatment with our leads ameliorated weight loss (Figures S2-6B; Table S5). Endoscopic analysis revealed a progressive increase in disease activity in all treatment groups receiving DSS. However, treatment with individual leads and MAP#4 consortium resulted in significantly lower disease activity, relative to the DSS only group on the final day of treatment (Figures S2-6C). Histological analysis of DSS-treated mice revealed extensive colon damage characterised by crypt loss, epithelial erosion, and ulceration (Figure 4B). Treatment with the individual leads or MAP#4 consortium resulted in a significant improvement in pathology characterised by crypt re-formation, re-epithelisation and reduced immune cell infiltration (Figures 4B-E, S2D-G, S3D-G, S4D-G, S5D-G, S6D-G). Other beneficial effects were noted across subsets of the four strains, including significant increases in goblet cells (MH21-1, MH23-1, MH32-1, MAP#4), mucin production (MH21-1, MH23-1), intraepithelial lymphocytes (MH21-1, MH23-1, MAP#4), and significant reductions in lipocalin-2, a biomarker of intestinal inflammation^36^ (MH23-1, MH27-1, MH32-1), relative to the DSS only control (Figures S2-3I, S5-6I, S2-3K, S2-3J, S6J, S3-5H). Compared to the DSS-treated mice, the prophylactic treatment with live strains did not result in significant differences in the alpha and beta diversity of the faecal microbiome at the last day of the experiment, suggesting a direct therapeutic effect of the leads on the host (Figures S7A-D).

**Figure 4.**
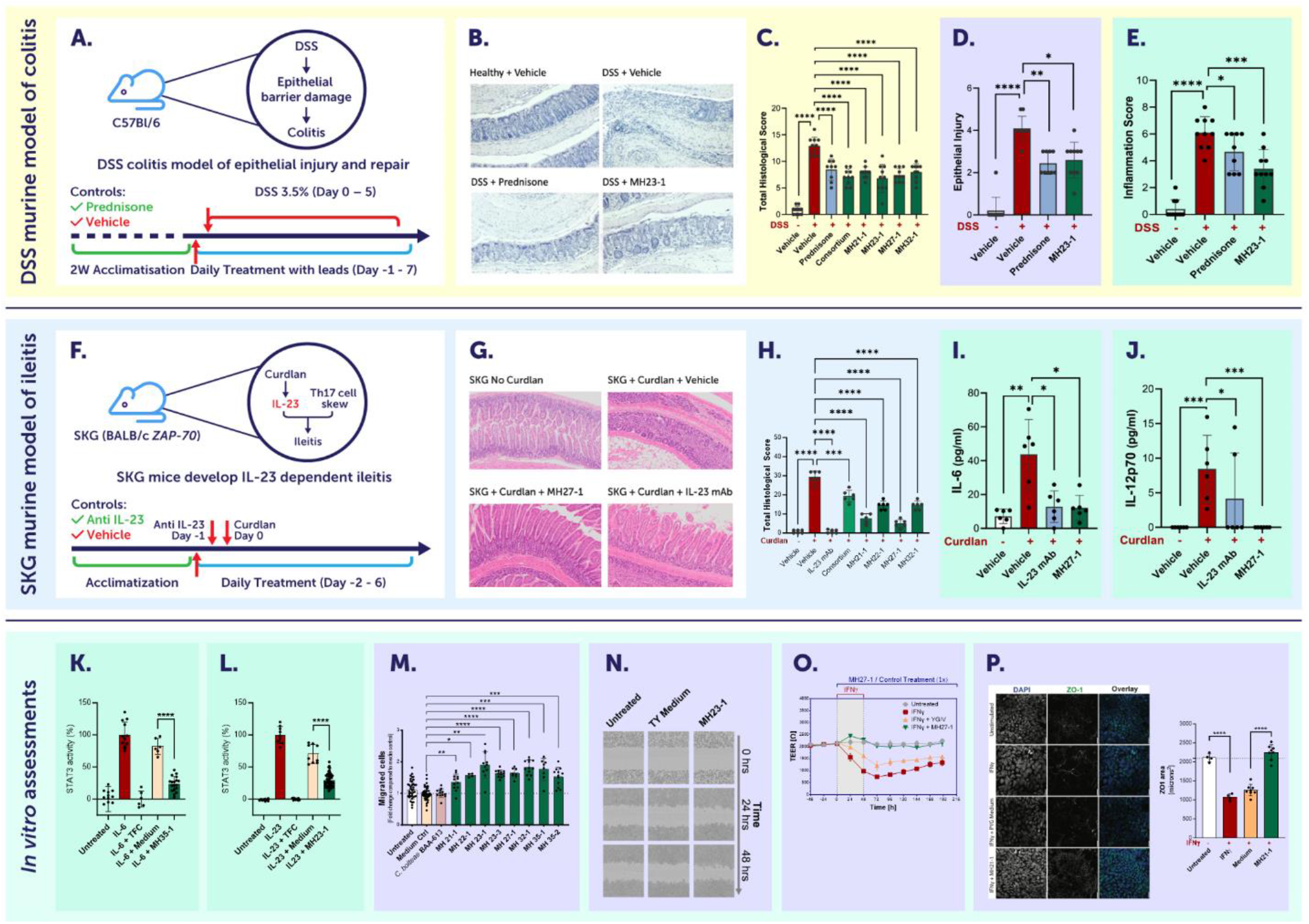
Exemplar data for assessing the potential of identified leads for therapeutic development. Activities relating to intestinal immunity and gut barrier integrity are shown in green and purple panels, respectively. Live cultures of MH21-1, MH23-1, MH27-1, MH32-1 and a consortium of these four leads (MAP#4) were assessed in a prophylactic dextran sulphate sodium (DSS) mouse model of colitis (n=10 per group) (A). Leads ameliorated DSS induced colitis as illustrated by representative gut histology images (B) and ameliorated increases in histopathological score, epithelial injury and inflammation compared to vehicle control (C^&^), (D^$^), (E^&^). Selected leads (MH21-1, MH22-1, MH27-1 and MH32-1, consortium MAP#4) were assessed in the SKG mouse model of ileitis (n=6 per group) (F). Leads ameliorated curdlan induced ileitis as illustrated by representative gut histology images (G) and ameliorated increases in histopathological score, IL-6 and IL-12p70 compared to vehicle control (H^&^), (I^&^), (J^$^). MH35-1 culture supernatant suppressed IL-6-induced STAT3 activation in HEKBlue IL-6 reporter cells (K^^^). MH23-1 culture supernatant suppressed IL-23-mediated STAT3 activation in HEKBlue IL-23 reporter cells (L^^^). Leads significantly increased the migration of HCT116 cells compared to medium control in a Transwell migration assay (M^^^). MH23-1 supernatant extract promoted faster wound closure compared to controls as illustrated using representative brightfield images of HCT116 cells in an IncuCyte scratch wound assay (N). T84 cells treated with MH21-1, MH23-1 and MH23-3 showed faster recovery from IFNγ mediated loss of barrier integrity, as assessed by transepithelial electrical resistance (TEER) (O^^^). Treatment with MH21-1 ameliorated IFNγ-induced reductions in ZO-1 expression in T84 cells (P^, scale bar 100 μm). For all data, ns: not significant; *, p < 0.05; **, p < 0.01; ***, p < 0.001; ****, p < 0.0001. All data presented as mean and standard deviation. ^$^Kruskal-Wallis test with uncorrected Dunn’s for multiple comparisons. ^&^Uncorrected Brown-Forsythe and Welch ANOVA test with multiple comparisons. ^^^T-test of medium control vs lead.

We also examined the efficacy of selected leads (MH21-1, MH23-1, MH23-3 and MH35-1) in a blinded therapeutic model of DSS-induced murine colitis. In this model, live leads were administered by oral gavage once daily for seven days at 2×10^8^ cells per mouse, starting on the fifth day of DSS treatment. In accordance with observations in the prophylactic model, treatment resulted in improvements in pathology including histological damage (all leads except MH23-1), epithelial injury (MH21-1), and reduced immune cell infiltration (all leads except MH23-1) (Figures S2M, S8-9B, S2O, S2N, S8-9C). Leads MH21-1 and MH23-3 also significantly reduced intestinal inflammation, as measured by lipocalin-2 (Figure S2P, S8E). At endpoint, DSS treatment resulted in a higher microbial beta diversity (Bray-Curtis dissimilarity to healthy mice) and reduced Shannon diversity, which was ameliorated by treatment with bacterial leads (Figures S7E-H).

We next tested the therapeutic activity of selected leads (MH21-1, MH22-1, MH27-1 and MH32-1) and the MAP#4 consortium in the SKG murine model of curdlan-induced ileitis (Figure 4F-J). SKG mice carry a mutation in the ZAP-70 gene and develop an IL-23 driven Crohn’s-like ileitis following disease initiation with curdlan treatment^37,38^. Leads were administered by oral gavage once daily for ten days at 2×10^8^ cells per mouse, starting two days prior to curdlan treatment. Histological analysis of curdlan-treated mice revealed significant damage to the ileum, characterised by infiltration of inflammatory cells and granuloma formation (Figure 4G). Treatment with an anti-IL-23 monoclonal antibody as positive control resulted in a significant improvement in histopathological score, as did treatment with the individual leads and MAP#4 consortium (Figure 4H), including reduced areas of inflammatory infiltrate (Figures S2U, S4-6N, S10C), granulomata (Figures S2V, S4-6O, S10D), and villous distortion (Figures S2W, S4-6P, S10E). There were also significant strain-specific reductions in pro-inflammatory serum cytokines, including IL-1α (MH22-1), IL-6 (MH21-1, MH27-1, MH32-1, MAP#4), IL-12 (MH21-1, MH22-1, MH27-1, MH32-1), IL-23 (MH21-1), TNF (MH21-1), and GM-CSF (MH21-1, MH22-1) (Figures 4I-J, S2Y, S4-6R, S10G). Notably, individual strains performed similarly to the MAP#4 consortium in the DSS and SKG models (Figures 4C, Figure 4H) suggesting that a combination treatment does not confer an additional benefit.

### Therapeutic leads modulate inflammatory pathways and promote wound healing

Given the beneficial anti-inflammatory effects observed *in vivo*, we next examined the ability of the bacterial leads to modulate IBD-associated immune pathways using human-derived reporter cell assays (Table 1). The IL-23-Th17 cell immune axis is central to the pathogenesis of IBD^39^. Therefore, we assessed if our leads can modulate the signalling pathways of IL-6 and IL-23 that are required for induction and maintenance of Th17 cells and regulation of their effector functions^39,40^. Specifically, we measured the effects on the transcription factor STAT3 (Signal Transducers and Activators of Transcription 3), which is activated by IL-6 and IL-23 and promotes excessive tissue inflammation and colitis in IBD^41^. Treatment of HEK-Blue-IL-6 reporter cells with cell-free supernatant prepared from cultures of all leads suppressed IL-6 mediated STAT3 activation relative to the medium control (Figure 4K, S2ZA, S3T, S4T, S5T, S8I, S9I). Similarly, culture supernatant from each lead, except MH27-1, suppressed IL-23 mediated STAT3 activation in HEK-Blue IL-23 reporter cells (Figure 4L, S2Z, S3S, S4S, S5S, S8H, S9H, S10H). The STAT3 suppressive effects were retained in <3 kDa fractionated supernatant indicating they were mediated by low molecular weight secreted bioactives.

**Table 1.**
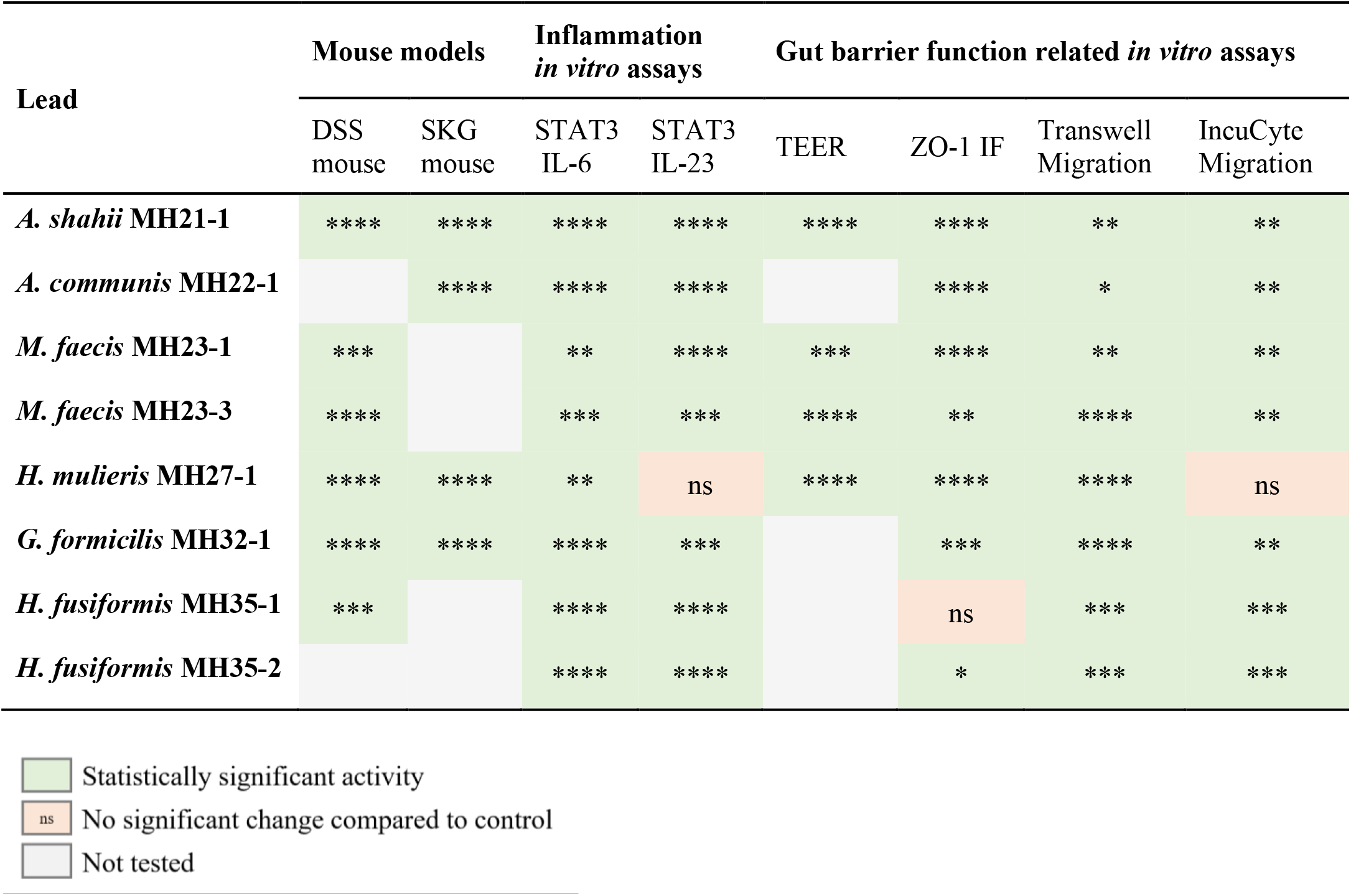
Overview of *in vitro* and *in vivo* assay results for eight microbial therapeutic leads. Indicated is whether treatment with the live therapeutic / bacterial culture supernatant vs the appropriate control (vehicle / medium control) resulted in a statistically significant effect, where ns: not significant; *, p < 0.05; **, p < 0.01; ***, p < 0.001 ****, p < 0.0001.

| Lead | Mouse models |  | Inflammation<br>*in vitro* assays |  | Gut barrier function related *in vitro* assays |  |  |  |
| --- | --- | --- | --- | --- | --- | --- | --- | --- |
|  | DSS mouse | SKG mouse | STAT3 IL-6 | STAT3 IL-23 | TEER | ZO-1 IF | Transwell Migration | IncuCyte Migration |
| *A. shahii* MH21-1 | **** | **** | **** | **** | **** | **** | ** | ** |
| *A. communis* MH22-1 |  | **** | **** | **** |  | **** | * | ** |
| *M. faecis* MH23-1 | *** |  | ** | **** | *** | **** | ** | ** |
| *M. faecis* MH23-3 | **** |  | *** | *** | **** | ** | **** | ** |
| *H. mulieris* MH27-1 | **** | **** | ** | ns | **** | **** | **** | ns |
| *G. formicilis* MH32-1 | **** | **** | **** | *** |  | *** | **** | ** |
| *H. fusiformis* MH35-1 | *** |  | **** | **** |  | ns | *** | *** |
| *H. fusiformis* MH35-2 |  |  | **** | **** |  | * | *** | *** |

**Table 1.** Overview of *in vitro* and *in vivo* assay results for eight microbial therapeutic leads.
|  |  |
| --- | --- |
|  | Statistically significant activity |
| ns | No significant change compared to control |
|  | Not tested |

Given the impact of our bacterial leads on the epithelial cell compartment and mucosal healing in the DSS and SKG mouse models, we examined their effects on gut epithelial cell function *in vitro*. Rapid migration of intestinal epithelial cells is a crucial component of wound healing to re-establish homeostasis after the gut barrier has been breached^42^. Treatment of human derived T84 gut epithelial cells with bacterial cell-free supernatant extract resulted in a significant increase in migration in a Transwell® migration assay for all leads (Table 1, Figures 4M, S2ZB, S3-5U, S8-10J). By contrast, cell migration was unaffected by treatment with the culture medium alone or the supernatant extract of *Enterocloster bolteae* BAA-613, a bacterium positively associated with IBD (Figure 4M). Similarly, we found a significantly accelerated rate of wound closure in the presence of the supernatant extract from seven of the eight tested bacterial leads, as assessed by scratch assay (Table 1, Figures 4N, S2ZC, S3-5V, S4W, S8-10K).

Gut barrier integrity was further evaluated using the trans-epithelial electrical resistance (TEER) assay with four of the leads. This assay uses T84 cells treated with the known barrier disruptor IFN-γ, which affects TEER by modulating expression of tight junction proteins (e.g. ZO-1, ZO-2, Occludin) thereby increasing paracellular permeability^43–45^. Treatment with IFN-γ resulted in an ∼80% reduction in TEER (Figure 4O), and subsequent treatment with supernatant extract from our leads resulted in faster recovery and higher gut barrier integrity relative to both the untreated and medium extract controls after twelve days (Table 1, Figures 4O, S2Zd, S3W, S4W, S8L). This finding was confirmed for MH27-1 using an additional readout for intestinal permeability, measuring the movement of fluorescent-labelled dextran across the gut epithelial cell monolayer at endpoint of the TEER assay (Figure S4X). We further showed that IFN-γ-induced reduction of ZO-1 protein expression in T84 cells was ameliorated by treatment with the supernatant extracts of all our leads, except MH35-1 (Table 1, Figures 4P, S3X, S4Y, S5W, S8M, S9-10L).

### Safety and manufacturability assessment

To explore the viability of our leads as LBP candidates, we used a combination of *in silico, in vitro* and *in vivo* studies to assess their safety and manufacturability. Genome analysis revealed that all leads were free of known human virulence factors (Table S6). No biosynthetic pathways for the production of harmful biogenic amines (i.e. histidine and tyramine) were detected, which can cause physiological symptoms when consumed in excess (Table S6). Using CARD and AMRfinderPlus, we detected homologs to several genes that confer resistance to antibiotics, but all matches were of low quality and were not co-located with mobile genetic elements that would facilitate horizontal gene transfer. Eight of the nine leads had homologs to genes conferring tetracycline resistance, consistent with the high prevalence of these genes in the gut microbiome^46^. However, only two of the leads grew in the presence of tetracycline (MH22-1 MIC 64μg/ml; MH23-1 MIC 16μg/ml). MH21-1 was also resistant to chloramphenicol, as predicted *in silico* and validated *in vitro*. All leads were sensitive to clindamycin when tested *in vitro*, despite two strains having low quality matches to genes conferring resistance to this antibiotic (Table S6). All leads were also sensitive to metronidazole and amoxicillin (except MH22-1 showed resistance to amoxicillin). All Bacillota (Firmicutes) strains were sensitive to vancomycin (Table S6). To further test safety and tolerability *in vivo*, we treated healthy mice with MH21-1, MH23-1, MH27-1 and MH32-1 for eight days and no changes in gut histology (e.g. total histological score, epithelial injury, inflammation score), colon length or colon weight were observed (data not shown). This was expected, given that the leads were isolated from healthy individuals and are commonly found in the healthy population. Similarly, no adverse effects of our leads were observed in the DSS or SKG experiments described above.

As a preliminary assessment of manufacturability, we tested our leads for phage activity as this can affect growth rate and cell yield. A prophage was identified in MH32-1 that showed high levels of spontaneous induction with lysis of the strain when treated with mitomycin C (Figure S11). The remaining leads did not show evidence for inducible prophages. We also assessed the ability of our top leads to grow on animal component free medium as this is an important consideration for LBP development. MH23-1, MH23-3, MH32-1 and MH27-1 grew well in YG/V and YG/P animal component free media, with growth of MH27-1 significantly improved relative to PYG, a nonselective medium widely used for the cultivation of anaerobic bacteria (Figure S12). By contrast, growth of MH21-1 in PYG or animal component free media was enhanced by the addition of haemin (Figure S12).

## Discussion

The microbiome therapeutics sector is gaining momentum with the first FDA approvals of faecal microbiota derived products to treat recurrent *Clostridioides difficile* infection^8,9^. These first products are based on faecal donor materials, which build on many years of efforts investigating the potential utility of faecal microbiome transplants. However, complexities with faecal donor material supply, variability and resulting efficacy necessitates the field to advance to rational development of therapies from defined and commercially manufacturable microbial strains and their therapeutic activities. To date, lead identification has largely relied on older technologies, including low-resolution 16S rRNA gene sequencing and standard isolation and screening approaches^13,14,47^, which are likely to miss the most relevant therapeutic leads and associated therapeutic biology. Here we demonstrate that the integration of high-resolution metagenomics combined with a well-powered disease cohort allows the discovery of potent therapeutic leads from the gut microbiome. Our extensive *in vivo* and *in vitro* data highlight the promise of this discovery approach and permitted the discovery of leads with potent activities. Notably, data guided discovery can identify keystone health-associated species with therapeutic potential from the thousands of microbial species that inhabit the human gut, thereby greatly expediting lead discovery. In combination with targeted isolation of desired lead bacteria, this approach substantially expands access to the gut microbiome, including members of the 63% of microbial species that so far have eluded cultivation^15^.

In this study, we demonstrated our data-driven approach for discovering novel microbiome derived therapies using IBD as a case study. Current IBD therapeutics, including corticosteroids, antimetabolites (e.g. methotrexate and thiopurines), small molecule inhibitors (e.g. JAK inhibitors) and monoclonal antibodies (e.g. anti-TNF, anti-IL-23, anti-integrin antibodies), primarily target the activated inflammatory response associated with the disease^48,49^. While these therapies are generally effective at suppressing inflammation, they are associated with potential adverse effects (e.g. weight gain, osteoporosis, liver or pancreas inflammation, more frequent infections, fever, nausea^48–50^) and many patients fail to achieve mucosal healing, which is a target of therapy and associated with improved long-term health outcomes. Microbiome derived therapies offer a new modality for treating IBD, with the potential for safe and effective long-term treatment^51^. To date, most efforts to harness the therapeutic potential of the microbiome for IBD have focused on conventional probiotic species (e.g. *Bifidobacterium, Lactobacillus* spp.), however the existing probiotic strains are ineffective in IBD^52,53^. More recent efforts have targeted fastidious species such as *F. prausnitzii*, which has been shown to possess anti-inflammatory properties^54,55^. Other microbiome targeting efforts for IBD treatment have focused on the development of consortia of human gut strains based on the assumption that these will be more therapeutically effective than single strains^56–58^. Notably, this was not the case in the present study – individual strains performed as well as a consortium of four leads (Figures 4C, Figure 4H) suggesting that a combination treatment provides no added benefit. Moreover, a single strain therapeutic would have significant chemistry manufacturing control (CMC) advantages, as the complexity of producing microbial consortia on a commercial-scale is highly challenging^59^. Single strain therapeutics are also advantageous from a regulatory perspective by simplifying manufacturing and stability, and demonstration of safety, efficacy and mechanism of action^60,61^.

We obtained isolates for seven of the twenty identified lead species, including three that were not described at the time of our analyses. To our knowledge, none of the lead species have previously been suggested as therapeutics for IBD. This highlights the potential of a precision metagenomics approach and importance of considering the entire microbiome when identifying LBP candidates. Encouragingly, treatment with our rationally identified leads significantly improved markers of disease histopathology, including crypt re-formation, re-epithelization and reduced immune cell infiltration, in both prophylactic and therapeutic models of IBD (DSS and SKG), and in the case of DSS models, treatment reversed endoscopically confirmed mucosal inflammation.

Serum levels of the pro-inflammatory cytokine IL-23 are higher in patients with IBD compared to healthy cohorts, and serum IL-23 levels correlate with severity of disease^62,63^. Similarly, increased levels of IL-6 in the colonic mucosa and blood of IBD patients has been observed and are associated with disease activity status^64^. IL-6 and IL-23 both activate STAT3 driving induction of acquired immune responses and survival of pathogenic T-cells, thereby resulting in excessive tissue inflammation and colitis^41,65–68^. Thus, agents that suppress STAT3 signalling hold great potential to attenuate gut inflammation and IBD severity. Our leads reduced serum cytokine concentrations in an SKG murine model of Crohn’s-like ileitis, including IL-6, IL-12, IL-23 and GM-CSF, suggesting they exert systemic effects. Lead strains also suppressed both IL-6 and IL-23 mediated STAT3 activation *in vitro*, suggesting this anti-inflammatory activity likely contributes to the improved disease markers observed in the *in vivo* IBD models.

Although anti-inflammatory agents can be effective for providing symptomatic relief of IBD, therapies that promote mucosal healing and restoration of gut barrier integrity are associated with improved long-term health outcomes and are emerging as new clinical targets^19,69,70^. We found that the secretome of our lead strains promoted cell migration and wound healing, and improved intestinal barrier integrity. The presented data indicate that several of our lead candidates may provide a dual therapeutic mechanism, suppressing key inflammatory pathways associated with IBD and promoting mucosal healing. This combined with the likely excellent clinical safety profile of LBPs demonstrates the potential of developing safe and efficacious therapeutics from commensal bacteria naturally occurring in the healthy human gut. Isolates from the lead species *A. shahii, H. mulieris*, and *M. faecis* had the overall best combination of efficacy and manufacturing potential. Strains from these three species have been assessed for manufacturability at a commercial manufacturer in preparation for human clinical trials.

In summary, we have identified new live biotherapeutic candidates from the healthy human gut microbiome that can regulate the immune system and improve gut barrier function, which show promise for effective long-term treatment of IBD. This demonstrates the potential of a data-driven approach for the rational design of well-defined microbial therapeutics with beneficial modes of action. We expect that this approach will be generalisable to any disease state with underlying microbiome aetiology and will expedite the identification and development of novel microbiome-derived therapeutics to improve human health.

### Taxonomic descriptions

#### Emended description of the genus *Hominicoprocola* Afrizal et al. 2023

The description remains as given by Afrizal *et al*. (2023)^49^ with the following emendations. The genus is classified within the family *Oscillospiraceae*, order ‘Oscillospirales’, class *Clostridia*, phylum *Bacillota* (Bacillota_A in GTDB 09-RS220). Cells are rod-shaped and approximately 10 μm in length or spindle-shaped cells with average 1–2 µm length on YCFA medium under anaerobic conditions.

#### Emended description of the species *Hominicoprocola fusiformis* Afrizal et al. 2023

The description remains as given by Afrizal *et al*. (2023)^49^ with the following emendations. The species features are based on description of three strains, including the type strain CLA-AA-H269^T^ (=DSM 113271^T^), MH35-1 (=originally deposited at the National Measurement Institute (Melbourne, Australia) as “*Colonithrix sana*” under accession number V21/019213) and strain MH35-2 (=originally deposited at the National Measurement Institute (Melbourne, Australia) as “*Colonithrix sana*” under accession number V21/019214). The two later strains were isolated from the stool specimen of a woman. It is predicted to use starch and produce acetate, propionate, l-glutamate and butyrate. It is also predicted to use glucose, fructose, gluconate, lactose, trehalose and lactaldehyde. It is predicted to be auxotrophic for alanine, tryptophan, histidine, lysine, methionine, phenylalanine and tyrosine. The genomic G + C content is 55.4 mol.% (contigs >5kb).

#### Description of *Vescimonas sanitatis* sp. nov (sa.ni.ta’tis. L. gen. n. *sanitatis* of health, referring to a healthy individual)

Cells are Gram-negative staining, spiral shaped and approximately 5 μm in length. It is predicted to use glucose, cellobiose, fructose and xylose. It is predicted to be auxotrophic for alanine, tryptophan, arginine, histidine, isoleucine, leucine, lysine, methionine, phenylalanine, tyrosine and valine. It is predicted to use lysine as an energy source. It is predicted to produce acetate and butyrate. The type strain MH37-1^T^ was isolated from the stool specimen of a woman. The genomic G + C content is 57.1 mol.% (contigs >5kb). The genome size is 2690112 bp.

#### Description of *Hominicoccus* gen. nov

*Hominicoccus* (Ho.mi.ni.coc’cus. L. masc. n. *homo* human being, man; N.L. masc. n. *coccus* a coccus; from Gr. masc. n. *kokkos* a grain or berry; N.L. masc. n. *Hominicoccus* a coccus from a human being)

The description of the genus is the same as for the type strain of *Ruminococcus gnavus* (Moore et al. 1976) comb. nov. Separation of the species into a new genus is justified by its distinct phylogenetic position on concatenated protein according to the GTDB within the family *Lachnospiraceae*, order *Lachnospirales*, class *Clostridia*, phylum *Bacillota* (Bacillota_A in GTDB 09-RS220). Type species: *Hominicoccus gnavus*.

#### Description of *Hominicoccus gnavus* comb. nov

*Hominicoccus gnavus* (gna’vus. L. masc. adj. *gnavus* busy, active [referring to the active fermentative ability of this species])

Basonym: *Ruminococcus gnavus* Moore et al. 1976 (Approved Lists 1980)

The description is the same as for *R. gnavus* given by Moore et al. 1976^91^. The type strain is ATCC 29149^T^ (= VPI C7-9 = JCM 6515).

#### Emended description of the genus *Hominenteromicrobium* Afrizal et al. 2023

The description remains as given by Afrizal *et al*. (2023) with the following emendations. The genus is classified within the family *Acutalibacteraceae*, order ‘Oscillospirales’, class *Clostridia*, phylum *Bacillota* (Bacillota_A in GTDB 09-RS220).

#### Emended description of the species *Hominenteromicrobium mulieris* Afrizal et al. 2023

The description remains as given by Afrizal *et al*. (2023) with the following emendations. The species features are based on description of two strains, including the type strain CLA-AA-H250^T^ (=DSM 113252^T^) and MH27-1 (deposited at the National Measurement Institute (Melbourne, Australia) under accession number V21/015888) isolated from the stool specimen of a man. Cells stain Gram-variable but are predominantly Gram-negative when grown in broth. It is predicted to use glucose, fructose, maltose, mannose, cellobiose, trehalose, raffinose, melibiose, lactose, galactose, sucrose, glucosamine, N-acetyl-D-glucosamine. It is predicted to be auxotrophic for alanine, tryptophan, phenylalanine, tyrosine, aspartic acid, glutamic acid, methionine, proline and serine. It produces acetate and succinate (data not shown) and is predicted to also produce formate and lactate. An antibiotic resistance gene related to tetracycline-resistance (*tetW*, ARO:0000002) was detected. The genomic G + C content of MH27-1 is 50.1 mol.% (contigs >5kb). The strain MH27-1 was originally classified and named as *Candidatus* Intestinicoccus colisanans by Zhou et al. 2023 based on the genomes sequence GCA_021029585.1, and its name validated under the SeqCode (seqco.de/r:tpd6ryk0). Based on priority dates, *Intestinicoccus colisanans*^Ts^ is a later synonym of *H. mulieris*.

## Acknowledgements

We thank Quinn Cao for assisting with the graphical design of the figures and the anonymous participants who contributed samples for this study. We thank Fabio Rigato for assisting with high performance cloud computing and Thomas Kryza for assistance with the IncuCyte system.

## Author contributions

LK, PÓC, PH, GWT conceived the original idea, and PÓC and LK planned the experiments and supervised the project. NA, DLAW, TM, LR, BW supervised components of the project. TK and SK supervised animal experiments. LK, PÓC, PH, AK, KT, JB drafted the manuscript. JB, AR, IHF, HC contributed to designing experiments and interpreting results; HC provided regulatory guidance. JB, DLAW, DHP, AA, PVW, MS, RN, PE, LK did bioinformatics analysis. JB, MN, MMC, LK did data analysis, machine learning and statistics. LK identified lead species and CV, BN, JJ, JZ, PÓC isolated the presented strains. AK, AR, JZ, BN, JKL, JJ, MB, KAM, LF, SMD, MC, DB, KZ, AP carried out wet lab experiments and/or contributed to the interpretation of the results. All authors discussed the results and contributed to the final manuscript.

## Data availability

Full access to the datasets presented in this manuscript will be provided upon reasonable request. Genomes of presented isolates will be provided by upload to a public repository and accession codes and unique identifiers will be made available prior to publication.

## Code availability

The custom code developed for this study can be made available upon reasonable request.

