## Supplementary material for "A rational approach for the targeted discovery and characterisation of microbiome-derived therapeutics": Methods and Supplementary Materials

***Australian cohort.*** Microba's Insight™ test provides an easy and comprehensive gut microbiome analysis using metagenomics. More than 4,065 Insight™ customers have consented to inclusion for research purposes. Based on this data, we have established a large human cohort with detailed medical history, lifestyle, phenotype and microbial metagenomic data for over 4,065 adults representing a cross section of the Australian population.

***Healthy control cohort.*** We have established a healthy cohort from an exclusive subset (n=241) of the Australian cohort based on strict health and lifestyle criteria. General inclusion criteria include: must be over the age of 18, BMI less than 30, no major medical conditions, no medications within the last 6 months that are known to significantly impact the gut microbiome (e.g. antibiotics, antifungals, antivirals, immunosuppressants, PPIs, statins, antidepressants, etc.), not a smoker, not pregnant, not exposed to hazardous chemicals, does not suffer from depression, anxiety or stress (DASS 21), eats more than 2 serves of fruits/veg daily, and less than 2 std alcoholic drinks per day.

This group consists of:

- 55% female, 45% male
- mean age of 47.2 (range 19 – 75) with a normal distribution
- a mean BMI of 23.3
- Ethnicity is predominately Australian/New Zealanders of European descent (78%), Western Europeans (8%), and Asian (4%)
- Education completed: University degree – 40.6%, College or professional certification – 26.2%, Master's degree or higher – 17.8%, Finished year 12 – 7.9%, Did not complete year 12 – 7.4%
- Relationship status: Married or in de facto relationship (opposite sex couple) – 78.7%, Single – 11.9%, Divorced – 3.5%, Married or in de facto relationship (same sex) – 3%

***Inflammatory Bowel Disease (IBD) cohort.*** From the 4,065 metagenomes obtained from a cross section of the Australian population, 80 individuals with self-reported diagnosis of IBD were selected to be included in the disease cohort. Of these, 30 subjects indicated a diagnosis of Crohn's disease (CD) and 50 subjects suffered from ulcerative colitis (UC). Exclusion criteria that were applied for the IBD cohort was an age filter (>=18 years), unknown gender, and missing specification for IBD subtype (CD or UC).

Population characteristics of this cohort were as follows:

- 58% female, 42% male
- mean age of 49.7 (range 21 – 88) with a normal distribution
- a mean BMI of 24.6
- Ethnicity is predominately Australian/New Zealanders of European descent (83.1%), Western Europeans (9.1%), and Asian (2.6%)
- Education completed: University degree – 40.3%, College or professional certification – 22.1%, Master's degree or higher – 15.6%, Finished year 12 – 6.5%, Did not complete year 12 – 13.0%
- Relationship status: Married or in de facto relationship (opposite sex couple) – 67.0%, Single – 26.0%, Divorced – 3.9%, Widowed – 1.3%

***Sample preparation***. Faecal samples were extracted using the DNeasy 96 PowerSoil Pro QIAcube HT Kit (Qiagen 47021) in 2 mL deep well plate format as per manufacturer's instructions with a modified initial processing step on the QIAcube HT DNA extraction system (Qiagen 9001793). Mechanical lysis was performed with PowerBead Pro beads (Qiagen 19311) and optimised volume adjustments enabling more efficient lysis of the complete microbial population. The kit contains streamlined inhibitor removal technology to eliminate the challenging inhibitors commonly found in stool samples. Resulting DNA was quantitated using a high sensitivity dsDNA fluorometric assay (QuantIT, ThermoFisher, Q33120). Samples were required to reach a minimum criteria of 0.2 ng/μL concentration to pass processing requirements.

***Metagenomic library preparation***. Libraries were constructed using the Illumina DNA Prep (M) Tagmentation Kit (Illumina, 20018705) with IDT for Illumina DNA/RNA UD Index Sets A-D (Illumina 20027213-16) according to manufacturer's instructions, with a modification of volume to accommodate processing in a 384 plate format. Resulting libraries were assessed using a high sensitivity dsDNA fluorometric assay (QuantIT, ThermoFisher, Q33120) and individual libraries were visualised with capillary gel electrophoresis using the QIAxcel DNA High Resolution Kit (Qiagen, 929002). Libraries were required to meet minimum criteria for average size, smallest and largest fragment gating and concentration.

***Metagenomic sequencing.*** Individual libraries were pooled in equimolar amounts to create a sequencing pool which was assessed using a high sensitivity dsDNA fluorometric assay (QuantIT, ThermoFisher, Q33120) and visualised with capillary gel electrophoresis using the QIAxcel DNA High Resolution Kit (Qiagen, 929002). Pools were required to meet minimum criteria for average size, smallest and largest fragment gating and concentration. Sequencing pools were loaded and sequenced on the NovaSeq6000 (Illumina) using v1.5 300 bp PE sequencing reagents, according to the manufacturer's instructions. Sequence data was reviewed for general minimum performance requirements for yield and sequence quality. The resulting data was additionally reviewed for performance requirements of known control samples that were included in each processing run and were required to meet minimum requirements for contribution of reads from reagents (background contamination) and for appropriate reporting of known control sample contents as assessed by Hellinger distance and associated statistical measures.

***Metagenomic sequencing data quality control (QC).*** Paired-end DNA sequencing data were demultiplexed and adaptor trimmed using Illumina BaseSpace Bcl2fastq2 (v2.20) accepting one mismatch in index sequences. Reads were then quality trimmed and residual adaptors removed using the software Trimmomatic v0.39[1] with the following parameters: -phred33 LEADING:3 TRAILING:3 SLIDINGWINDOW:4:15 CROP:100000 HEADCROP:0 MINLEN:100. Human DNA was identified and removed by aligning reads to the human genome reference assembly 38 (GRCh38.p12, GCF_000001405) using bwa-mem v0.7.17[2] with default parameters except minimum seed length set to 31 (-k 31). Human genome alignments were filtered using SAMtools v1.7, with flags -ubh -f1 -F2304. Any read pairs where at least one read mapped to the human genome with >95% identity over >90% of the read length were flagged as human DNA and removed. All samples were then randomly subsampled to a standard depth of [seven million] read pairs.

***Reference microbial genome database.*** We have established an extensive, high quality genome database – the Microba Genome Database (MGDB) – which is used as a reference for the identification of microbial species and functions from metagenomic sequence data. The database covers all prokaryote lineages but has been specifically enriched for genomes mined from faecal metagenomes. The MGDB has been compiled from publicly available microbial genomes and metagenome-assembled genomes (MAGs) that we mined from Insight$^{TM}$ samples and publicly available datasets. All included microbial genomes have been quality controlled through rigorous standards, clustered into species groups, and systematically functionally annotated. Species represented in the MGDB have been systematically named using an in-house version of the Genome Taxonomy Database (GTDB)[3]. In detail, the MGDB comprises 400,971 high quality prokaryotic genomes from the following sources: the Genome Taxonomy Database (GTDB, including GenBank and RefSeq genomes), metagenome-assembled genomes (MAGs) mined from Insight$^{TM}$ samples, MAGs mined from sequence read archive (SRA) samples, and MAGs and isolate genomes from the Almeida et al[4], Forster et al[5], Nayfach et al[6], Pasolli et al[7], and Zou et al[8] studies. All genomes were quality controlled, with an average completeness of 92.43% and average contamination of 1.15% as estimated with CheckM[9]. Genomes were clustered into Operational Species Clusters and representative genomes were selected from these clusters such that the phylogenetic diversity within a species cluster is maximised. The resulting MGDB v2 database contained 73,646 representative genomes grouped into 28,246 species clusters.

***Quantification of microbial species abundances.*** Species profiles were obtained with the MCP v2.0[10] using the MGDB v2 as the genome reference database. The community profiles produced by the MCP were based on the rank normalised taxa and comprehensive species clusters defined by the Genome Taxonomy Database[3].

***US cohort.*** In our meta-analysis we included faecal metagenomic data from a US based IBD cohort (Harvard). Previously published metagenomic sequence data[11] was downloaded from SRA and re-analysed using the MCP, as described above.

***Cohort matching.*** Healthy control and IBD samples from the Australian cohort were matched for potential confounding factors. Individuals >18 years old and with completed age and gender data were included. Healthy individuals and individuals with IBD, UC and CD were age- and gender-matched by propensity score matching using the MatchIt v4.4.0 package in R[12]. The ratio of control to disease group size was selected as the highest ratio at which the total standard mean difference (age) and unstandardised difference in proportion (gender) was <0.1.

***Meta-analysis.*** MCP community abundances for the US and matched Australian cohort were merged for meta-analysis. For each condition (IBD, CD, UC), species prevalences within the healthy and disease cohorts were compared using the Cochran–Mantel–Haenszel test (CMH) using the dataset of origin as a confounder (US, Australian cohort). P-values resulting from this analysis were corrected for multiple testing using FDR. The top 20 healthy-associated species were selected for each condition by ranking the FDR corrected CMH p-values, omitting species that were 1) in >55% of diseased samples, 2) in <30% of healthy samples, and 3) disease prevalence was greater than healthy prevalence.

***Diversity analysis.*** Sample Shannon diversity, evenness and richness was calculated for samples in the Australian cohort using vegan (v2.6-4)[13]. Diversity, evenness and richness was compared between healthy, IBD, UC and CD using a t-test. P-values were corrected for multiple testing using FDR.

***Machine learning-based lead discovery.*** Machine learning was used to identify microbial species important for differentiating microbial community profiles from controls versus individuals with IBD, using species profiles from the Australian and US-based cohorts. Relative abundance data were pre-processed using recipes v1.0.6[14], including square root transformation and removal of features with zero variance. Deep Learning Neural Networks were fit using keras v2.9.0[15], tensorflow v2.9.0[16] and tune v1.1.1[17]. Gradient Boosting Machines (GBM) were fit using H2O v3.38.0.1[18]. In both cases, hyperparameter tuning and performance estimation were performed using resampling with 5-fold cross validation, with identical folds calculated using rsample version 1.1.1[19]. H2O 3.38.0[18] was used to calculate species contributions to model predictions with Shapley Additive exPlanations (SHAP) values[20] and variable importance.

***Metabolic reconstruction.*** Protein coding sequences were predicted and annotated using the annotate function in enrichM (0.6.2). Briefly, enrichM identifies protein coding sequences using prodigal (version) in -p meta mode. The amino acid sequences were then annotated using emapper, providing E.C., TCDB and eggnog classifications[21,22]. HMMER hmmsearches (version 3.1b2) against Pfam (release 33.0), TIGRFAM (release 15.0) and dbCAN2 (downloaded September 2019) were used to annotate functional domains, key metabolic markers and carbohydrate activate (CAZy) enzymes, respectively. Metabolic pathways were identified using the classify function in enrichM, which assesses annotations and their genomic position against manually defined metabolic pathway definitions. A pathway was considered present in a genome if it encoded >80% of the required proteins and passed all required synteny checks. These automatically predicted pathways were then manually assessed. In addition, gutSMASH (version 1.0.0)[23] was applied to identify common functions mediated by gut microbiomes.

***Bacterial strains, culture conditions and analyses.*** Stool samples were collected from healthy adults (see 'Healthy control cohort' for detailed inclusion criteria) following informed consent and the ethical guidelines approved by Bellberry Limited (HREC2018-05-400-A-6). The stool samples were mixed with an equal volume of sterile oxygen free buffered glycerol solution in a Coy vinyl anaerobic chamber (85% $N_2$:10% $CO_2$:5% $H_2$ atmosphere) and then stored at -80°C until required. The bacterial strains described in this study were processed in a Coy vinyl anaerobic chamber. All strains were routinely grown in TY (Tryptone 10 g/L, yeast extract 2.5 g/L, glucose 4 g/L, cellobiose 1 g/L, maltose 1 g/L, haemin (500 mg/L) 10 mL/L, acetic acid 1.9 mL/L, salt solution 2[24] 38 mL/L, salt solution 3[24] 38 mL/L, sodium bicarbonate 8 g/L, resazurin (0.1% w/v) 1mL/L, L-cysteine 1 g/L) or PYG (Tryptone 20 g/L, yeast extract 10 g/L, glucose 10 g/L, haemin (500 mg/L) 1 mL/L, salt solution 2[24] 38 mL/L, salt solution 3[24] 38 mL/L, sodium bicarbonate 8 g/L, resazurin (0.1% w/v) 1mL/L, L-cysteine 1 g/L) based media unless otherwise stated. All strains were identified by whole genome sequencing and stocked by mixing actively growing culture with an equal volume of glycerol solution and storing at −80 °C. Isolates were identified using the MGDB ANI pipeline, which uses MASH to generate a list of MGDB representative genomes that have >90% ANI with the target genome. The closest match is then identified using fastANI[25,26] and species level taxonomy is inherited if the match is >95% ANI and > 0.65 alignment fraction. The purity of the stocked isolates was assessed by microscopy and analysis of the whole genome sequence using CheckM[9]. For CheckM, if the contamination was >0% the genome and corresponding unassembled sequences were manually inspected for contamination by comparative genomics and by mapping against the MGDB. Growth optimisation experiments were performed using a Multiskan Go 100-240V (MH23-1, MH23-3, MH27-1, MH32-1) and Cerillo Stratus (MH21-1) plate readers. Specific growth rates were calculated as previously described[27].

***Isolation of lead candidates.*** Isolates of *A. shahii*, *A. communis*, *M. faecis*, *G. formicilis* were produced using traditional culturing approaches. Briefly, *A. shahii* MH21-1 was isolated by inoculating a donor faecal sample with *A. shahii* present at a relative abundance of 0.24% into M4 broth (NZ-amine 10 g/L, yeast extract 2.5 g/L, L-arabinose 2 g/L, clarified rumen fluid 30 ml/L, salt solution 2[24] 38 mL/L, salt solution 3[24] 38 mL/L, sodium bicarbonate 8 g/L, sodium azide (10% w/v) 2.5 mL/L, resazurin (0.1% w/v) 1mL/L, L-cysteine 1 g/L). The enrichment culture was subsequently plated for colonies on *Bacteroides* Bile Esculin Agar. Single colonies were picked and by this approach and *A. shahii* MH21-1 was identified following whole genome sequencing. *A. communis* MH22-1 was isolated by plating a donor faecal sample with *A. communis* present at a relative abundance of 1.55% onto *Bacteroides* Bile Esculin Agar. Single colonies were picked by this approach and *A. communis* MH22-1 was identified following whole genome sequencing. *M. faecis* MH23-1 was isolated by inoculating a donor faecal sample with *M. faecis* present at a relative abundance of 0.47% into Schaedler broth and then serially diluting to extinction. The dilution-to-extinction culture series was sequenced and an enrichment culture with *M. faecis* at 14% relative abundance was identified. This enrichment was subsequently diluted to extinction again in Schaedler broth and an enrichment with *M. faecis* at 42% relative abundance was identified. Colonies were recovered by streaking on *Bacteroides* Bile Esculin Agar with the aminoglycoside antibiotics omitted and an isolate termed *M. faecis* MH23-1 was identified by whole genome sequencing. *M. faecis* MH23-3 was produced by inoculating a donor faecal sample with *M. faecis* at 0.12% relative abundance into Yeast *N*-acetylglucosamine broth (Yeast extract 10 g/L, *N*-acetylglucosamine 10 g/L, salt solution 2[24] 40 mL/L, salt solution 3[24] 40 mL/L, sodium bicarbonate 8 g/L, resazurin (0.1% w/v) 1 mL/L, L-cysteine 1 g/L) and then serially diluting to extinction. An enrichment with *M. faecis* at 47.8% was identified and colonies were subsequently produced by streaking on TY medium supplemented with 0.5% w/v sodium azide. Single colonies were picked and *M. faecis* MH23-3 was identified by whole genome sequencing. *G. formicilis* MH32-1 was isolated by inoculating a donor faecal sample with *G. formicilis* present at a relative abundance of 2.48% into TY medium and then serially diluting to extinction. Colonies were recovered on TY medium and then screened as previously described for extremely oxygen sensitive isolates[28]. The extremely oxygen sensitive isolate *G. formicilis* MH32-1 was identified by whole genome sequencing.

Isolates of *H. mulieris*, *H. fusiformis* and *V. sanitatis* were produced by genome directed isolation[29]. Briefly, the metabolic pathways of for *H. mulieris*, *H. fusiformis* and *V. sanitatis* were reconstructed and custom media designed for each species to fulfil their predicted nutritional requirements (see Metabolic reconstruction). Briefly, potential donor samples were mined for MAGs of the target species, which were analysed together with high-quality representatives from the MGDB (>90% complete, <5% contamination) to create a sample-specific genome database to infer nutritional requirements (see Metabolic reconstruction). Energy sources were derived from carbohydrate (fibres, oligo- and monosaccharides), amino acid degradation (Stickland reactions), and fermentation pathways that were annotated in the target genome database. Candidate energy sources were ranked by the energy produced (carbohydrates preferred), whether the pathway is predicted as highly expressed (average CAI), and how selective the energy source is for the target species compared to the background community. Amino acid requirements were inferred from auxotrophies in the target species. Any amino acid pathway that was absent in the target genomes was added to the media. While vitamin requirements were assessed in the standard pipeline, most nutritional requirements were fulfilled for *H. mulieris, H. fusiformis* and *V. sanitatis* through the addition of a complete vitamin solution (see Supplementary Table S1 for vitamin solution (1000x) recipe). *H. mulieris* MH27-1 was isolated by inoculating a donor faecal sample with *H. mulieris* present at a relative abundance of 0.37% into custom medium (see Supplementary Table S1) and then serially diluting to extinction. The dilution-to-extinction culture series was sequenced and a low diversity enrichment culture with *H. mulieris* at a relative abundance of 77% was identified.

This enrichment culture was then used to establish a further dilution-to-extinction culture series and a second low diversity enrichment with *H. mulieris* at over 79% was identified. Colonies were recovered on PYG agar and subsequently picked into PYG broth. From this, a bi-culture comprised of *H. mulieris* and *Ruthenibacterium lactatiformans* was identified. To produce an axenic culture of *H. mulieris*, the bi-culture was transferred into YCFAmod (NZ-amine 2 g/L, yeast extract 0.5 g/L, soluble starch 0.5 g/L, glucose 0.5 g/L, cellobiose 0.5 g/L, haemin (500 mg/L) 20 mL/L, salt solution 2[24] 38 mL/L, salt solution 3[24] 38 mL/L, volatile fatty acid mix (propionate (9 mM), isobutyrate, (1 mM), isovalerate (1 mM), valerate (1 mM)) 3 mL/L, sodium bicarbonate 8 g/L, resazurin (0.1% solution) 1 mL/L, L-cysteine 1 g/L, vitamin solution (1000x) 1mL/L) broth and the enrichment culture was then streaked onto YCFAmod agar. Two distinct colony types were observed after several days, and by this approach *H. mulieris* MH27-1 was identified following whole genome sequencing. *H. fusiformis* MH35-1 and MH35-2 were isolated by inoculating a donor faecal sample with *H. fusiformis* present at a relative abundance of 1.2% into custom broth (see Supplementary Table S1) and then serially diluting to extinction. The donor sample also contained a related species ER4 sp900317525 at a relative abundance of 0.04% and xylose was also added to the custom medium to support growth of this bacterium. The amino acids in the custom broth fulfilled ER4 sp900317525 auxotrophies and acted as a carbon source for *H. fusiformis*, which is predicted to ferment serine to acetate or further to butyrate. The dilution-to-extinction culture series was sequenced and a low diversity enrichment culture with *H. fusiformis* at a relative abundance of 6.6% was identified. Colonies were recovered and screened by PCR using *H. fusiformis* specific primers (Pf 5′ ATC GGT GGT CAT GAT GGC GTA GGC AGC C; Pr 5′ AAG ACG CCC GCA CCG TTG AGC GCG AGG). By this approach, *C. sana* MH35-1 and *H. fusiformis* MH35-2 were identified following whole genome sequencing. *V. sanitatis* MH37-1 was isolated by inoculating a donor faecal sample with *V. sanitatis* present at a relative abundance of 0.86% into custom broth (see Supplementary Table S1) and then serially diluting to extinction. The dilution-to-extinction culture series was sequenced and a low diversity enrichment culture with *V. sanitatis* at a relative abundance of 31% was identified. Filter sterilised cell free culture supernatant was prepared from this enrichment and a further dilution to extinction enrichment was established using the *V. sanitatis* enrichment as an inoculum. The dilution-to-extinction enrichments were spotted in 10 µL volumes onto a TY plate. Colonies were patched and screened by PCR using *V. sanitatis* specific primers (Pf 5′ GAA CCG TGT CTC GCC GCC CCG GCG; Pr 5′ CTG CCG TGC CTT CGG GCC GGA CCG). A PCR positive patch was identified and shown by sequencing to be a low diversity enrichment comprised of MH37 and several other species. The enrichment was serially diluted and plated onto TY agar. Colonies were screened and by this approach *V. sanitatis* MH37-was identified by whole genome sequencing.

***In-silico* safety analysis.** Protein coding sequences from the isolate genomes were assessed for the presence of antimicrobial resistance (AMR) genes by blasting against the CARD database[30] (downloaded June 2021) using diamond blastp (50% AAI, 80% coverage) and AMRFinderPlus[31]. CARD hits were sorted into 'strict' or 'loose' based on whether the bit score was above or below the curated CARD cut-offs. Significant hits from both approaches were further annotated with CD-SEARCH to verify the annotations. Protein sequences were also searched against the Virulence Factor Database[32] (downloaded August 2021) using the same filters as above. Hits were further annotated with CD-SEARCH to verify the annotations and manually assessed.

Each lead was checked for the presence of key genes involved in the biosynthesis of biogenic amines, including clinically significant compounds such as histamine and tyramine. E.C.

annotations of the protein coding genes from Eggnog-mapper were provided to enrichM, along with custom pathway definitions for each biogenic amine (Supplementary Table S5). A pathway was considered complete if all enzymes were present and was considered partially complete if key enzymes were present and one or more other enzymes were missing from the pathway.

Isolate genome sequences were searched for mobile genetic elements (MGEs) using MobileElementFinder under default conditions[33]. Further, oriTfinder[34] was used to identify known AMR genes, virulence factors, conjugation machinery and other features associated with the replication of mobile genetic elements such as the origin of transfer (*oriT*). Plasmids were identified using PlasForest with default settings[35]. Regions designated as potential mobile genetic elements were checked for the presence of AMR and virulence genes.

***Phylogenetic analysis.*** A genome tree was constructed using representative genomes for each species in the MGDB using GTDB-Tk v1.7.0[36] with default settings. GTDB-Tk initially calls protein coding genes within MGDB genomes using prodigal (v2.6.3) and identifies 120 conserved single copy marker genes identified using Pfam, and TIGRFAM HMMs. These proteins are then aligned to their respective HMMs and merged into a single multi sequence alignment (MSA). The MSA is filtered to remove highly variable and conserved positions and subsampled to ~5,000 columns to reduce computational requirements. A phylogeny is then inferred using FastTree (v2.1.11) under the WAG+GAMMA models[37]. Non-parametric bootstrap values were calculated with 100 replications.

***Preparation of bacterial strains for animal experimentation***. Bacterial strains were grown to early stationary phase and the cell density of each culture was determined using a Helber Counting Chamber. The individual cultures were then centrifuged under a layer of sterile heavy mineral oil at 5,000 g for 10 minutes. The cell-free supernatant was discarded and cell pellets washed with 1.5 mL of sterile anaerobic buffered diluent (salt solutions 2 & 3[24], 38 mL/L each, resazurin (0.1% w/v solution) 1 mL/L, L-cysteine 1 g/L) and then centrifuged again. Finally, the washed cell pellet was resuspended in half strength glycerol solution (15% v/v glycerol solution in anaerobic buffered diluent) to a final concentration of 1 x $10^9$ cells/mL, aliquoted and frozen at -80°C until required. The MAP#4 consortium was prepared by mixing equal volumes of 1 x $10^9$ cells/mL of *A. shahii* MH21-1, *M. faecis* MH23-3, *H. mulieris* MH27-1 and *G. formicilis* MH32-1 strain to produce a cell consortium with each strain present at 2.5 x $10^8$ cells/mL and with a total cell count of 1 x $10^9$ cells/mL. The viability of the cell preparations was confirmed by thawing a single aliquot and streaking on an agar plate. The identity and purity of the individual strain preparations was confirmed by whole genome sequencing.

***Acute model of DSS induced colitis.*** Six-week-old C57BL/6 female mice purchased from Animal Resources Centres (Western Australia, Australia) or Australian Bioresources (NSW, Australia) were randomised and then co-housed for seven days prior to experimentation. To induce disease in the prophylactic model, mice were treated with 3.5% DSS *ad libitum* in the drinking water for 6 days. Naïve age matched control mice were processed and received DSS free drinking water. Prophylactic treatments started one day prior to provision of DSS and all mice were sacrificed 2 days after the final DSS treatment. For the treatments, mice were anesthetised with isofluorane and orally gavaged with 200 µL of bacterial preparations or vehicle control. Prednisone (2 $mg.kg^{-1}$) was administered following anesthetisation by intraperitoneal (i.p.) injection. Body weights and stool consistency were recorded daily. Stool samples were collected daily. Following sacrifice, the colon, liver and spleen were collected for analysis. Blood was collected by cardiac puncture. The therapeutic model was similarly performed except that the animals were treated with 2.5% DSS for a total of 5 days. The treatments commenced on final day of DSS administration, and all mice were sacrificed 6 days later. Prophylactic and therapeutic treatments were blinded during treatment phases and unblinded after data analysis. The DSS animal experiments were performed with ethical approval by the University of Newcastle Animal Ethics Committee (A-2020-021). In both DSS models, a minimum of ten animals per experimental group were used.

***Endoscopic and Histological scoring of DSS colitis.*** Animals were examined with a small animal gastrointestinal endoscope (Karl Storz Endoskope, Tuttlingen, Germany) on days -1, 2 and 6 for the prophylactic model, and days -1, 3 and 9 for the therapeutic model, to assess the extent of colon mucosal inflammation[38,39]. Briefly, mice were anesthetised with isoflurane and a colonoscope was inserted through the rectum. Images captured by high-definition videos were examined in a blinded manner to assess the presence and extent of disease pathologies (Supplementary Table S2). Histological scoring was performed essentially as described by Marks et al[38]. Briefly, samples were fixed in 4% formalin, paraffin embedded and sectioned. Tissue sections were hematoxylin and eosin stained to assess disease pathology and with Alcian blue to assess mucin production. Slides were imaged using the Aperio digital imaging system (Leica Biosystems, Nußloch, Germany). To grade colitis severity, the extent of inflammation and epithelial injury in the tissue sections were graded semi-quantitatively using an established scoring system (Supplementary Table S3). The samples were then randomised and subsequently scored in a blinded manner by a trained gastrointestinal pathologist.

***Mouse faecal metagenome analysis.*** Mouse faecal samples were collected on day -2, 0 and 6 in the prophylactic DSS experiment, and on day 7 and 10 in the therapeutic experiment. Only fresh faecal pellets were collected, immediately frozen on dry ice and then stored at -80°C until further processing. DNA extraction, library preparation and sequencing was performed as described above for human faecal samples. Faecal pellet metagenomes were processed using the MCP to determine community structure. PCAs were generated from species read counts that were hellinger transformed and scaled. Binary bray-curtis distances were calculated between untreated and treated samples using the vegdist from the vegan package. Shannon diversity, richness and evenness were determined from the hellinger transformed read counts using vegan. Bray-curtis distances, diversity, richness and evenness were compared between untreated and treated samples using Wilcoxon test. P-values were corrected for multiple testing using FDR.

***SKG model of murine ileitis.*** Five-week-old SKG mice were randomised and then co-housed for 3 weeks prior to experimentation. To induce disease, mice were administered curdlan (1,3-β-glucan, 3 mg per mouse) i.p.. Naïve age matched control mice were similarly processed except that they were administered saline i.p.. All treatments (n=6 per group) started prior to administration of curdlan, and all mice were sacrificed 7 days after the curdlan treatment. For the bacterial treatments, mice were orally gavaged with 200 μL of bacterial preparations or vehicle control starting 2 days prior to the curdlan administration. Anti-mouse IL-23 p19 monoclonal antibody (Thermo Fisher, 30 μg per mouse) was administered by i.p. injection one day prior to curdlan administration. Stool samples were collected daily. Body weights were measured at Day -2, 0 and 7. Following sacrifice, the colon, distal small intestine, mesenteric lymph nodes and spleen were collected for analysis. Blood was collected by cardiac puncture. The SKG experiment was performed with ethical approval by the University of Queensland Animal Ethics Committee (AEC QUT/TRI362/20).

***Clinical and histological scoring of SKG ileitis.*** Histological scoring was performed essentially as described by Benham et al[40]. Briefly, samples were fixed in 4% formalin, paraffin embedded and sectioned. Tissue sections were hematoxylin and eosin stained to assess disease pathology. Slides were imaged using an Olympus VS120 slide scanner (Olympus Corporation, Tokyo, Japan). To grade ileitis severity, the extent of inflammation and epithelial injury in the tissue sections were graded semi-quantitatively using an established scoring system (Supplementary table S4, Benham et al[40]). The samples were then randomised and subsequently scored in a blinded manner by a trained pathologist.

***Quantification of SKG serum cytokines.*** To quantify serum cytokines, the mice were euthanised by $CO_2$ asphyxiation after which blood was collected by cardiac bleed and serum isolated by centrifugation. Serum samples were stored at -80°C until quantitation. Serum cytokines were quantified using the LEGENDplex Mouse Inflammation Panel (13-plex) (BioLegend) according to the manufacturer's instructions to determine effects on cytokine production (IL-23, IL-1 α, IFN-γ, TNF, MCP-1, IL-12p70, IL-1β, IL-10, IL-6, IL-27, IL-17A, IFN-β, GM-CSF).

***Preparation of bacterial supernatants and extracts for in vitro assays.*** To prepare bacterial supernatant samples for *in vitro* assays, three independent colonies were inoculated and grown until early stationary phase thereby generating three seed culture broths. Then, each seed culture broth was used to inoculate two technical replicates each thus generating six technical replicates. The replicates were grown until early stationary phase and then cell free culture supernatant was harvested as previously described[41]. Culture supernatants were size fractionated by passing through a 3 kDa *CENTRICON*® column according to the manufacturer's (Merck Millipore) instructions.

Bacterial supernatant extracts were prepared using an Amberlite XAD-7 resin essentially as described by Colosimo et al[42]. Briefly, three single colonies of the bacterium of interest were inoculated and grown until early stationary phase. Each seed culture broth was then used to inoculate 600 mL of broth and the culture was again incubated until early stationary phase. Culture supernatants were prepared by centrifuging the culture at 4000 g for 30 minutes and then passing the cell free supernatant through a 3 kDa filter according to the manufacturer's instructions (Sartorius Vivaflow® 50 Ultrafiltration Unit 3kDa MWCO PES). Activated Amberlite XAD-7 resin was added to 400 mL of 3 kDa filtered cell-free supernatant (10% w/v), and the slurry was gently shaken overnight at 4°C. The resin was collected, washed with 400 mL of deionised water and then mixed with 120 mL of 100% methanol. Following 2 h incubation with gentle shaking, the methanol elution was collected. A second elution in 120 mL of 100% methanol was performed as previously described and the two elutions were ultimately combined and dried under vacuum using a rotary evaporator. The extract was fully resuspended in 100% DMSO and stored at -20°C.

***Quantitation of reporter gene activity.*** The ability of bacterial leads to modulate IL-6 mediated STAT3 activity in human cells *in vitro* was assessed using the HEK-Blue™ IL-6 reporter cell line (Invivogen). The HEK-Blue™ IL-6 reporter cell lines are stably transfected with the IL-6 receptor, STAT3 and a Secreted Embryonic Alkaline Phosphatase (SEAP) reporter gene. Stimulation with IL-6 causes STAT3-dependent expression of SEAP which is secreted into the medium and can be quantified using QUANTI-Blue solution (Invivogen). HEK-Blue™ cells were cultured in Dulbecco´s Modified Eagle Medium (DMEM) supplemented with 10% FBS. For the IL-6 assays, cells were seeded in 96-well plates (50,000 cells per well) and the following day, bacterial culture supernatant or sterile bacterial medium were added to cells at a final concentration of 5-10% v/v. After a 60-minute pretreatment, recombinant human IL-6 (2 ng/mL; R&D systems) was added. Additionally, IL-6 stimulated cells were simultaneously treated with the Janus kinase (JAK)-STAT inhibitor tofacitinib (10 μM) as negative control. After incubating the plates at 37°C for 24 hours, the STAT3 regulated SEAP reporter activity was assessed using QUANTI-Blue solution as recommended by the manufacturer. To quantify SEAP levels, the optical density of each well was read using a PHERAstar FS plate reader (BMG Labtech) at 630 nm. Results are the average of three independent experiments, and each condition had at minimum three technical replicates in each experiment. In addition to the measurement of STAT3 activity, potential cytotoxic effects of the treatments was assessed by using the CellTiter-Glo® 2.0 Cell Viability Assay as recommended by the manufacturer (Promega, Australia). The HEK Blue IL-23 reporter cell assay was performed similarly except that the cells were co-treated with IL-23 (5 ng/mL final concentration) and bacterial supernatants (10-25% v/v final concentration) as appropriate. IL-23 stimulated cells were also treated with tofacitinib (10 µM) as negative control. Treated cells were incubated at 37°C and SEAP activity was assessed after 6 hours. Statistical analysis of differences in SEAP activity was performed using the GraphPad Prism 9 software.

***Measurement of TEER activity.*** The impact of bacteria on gut barrier integrity was assessed by measuring the transepithelial electrical resistance (TEER) across confluent monolayers of T84 gut epithelial cells. T84 cells were obtained from Cell Bank Australia and cultured in DMEM Nutrient Mixture 12 (DMEM/F12; ThermoFisher Scientific, Waltham, MA, USA) supplemented with 5% foetal bovine serum and 1% Penicillin-Streptomycin. T84 cells were seeded into 24-well Millicell® polycarbonate cell culture inserts with 0.4 µm pore size (PSHT010R5) at a density of 60,000 cells per well in 400 µL medium, and 24 mL of medium was added into the single-well feeder tray. Medium changes of both compartments were performed every second day. After seven days of culture, the upper part of the plate assembly with cell culture inserts was transferred from the feeder tray to a 24-well receiver tray (PSMW010R5), with each receiver well containing 800 µL cell culture medium. TEER values of each well were measured daily using the Millicell® ERS-2 Voltohmmeter. The experiment was started once all cell monolayers reached stable TEER readings above 1500 Ω. To disrupt barrier integrity, IFNγ (50 ng/mL) was added to the basolateral compartment for 48 hours and then removed. To test if bacterial leads can enhance the recovery from IFNγ-mediated barrier function loss, 10% v/v 3kDa-filtered bacterial culture supernatant or the appropriate 3kDa-filtered bacterial medium control, were each diluted in T84 medium and added to the apical compartment. The DMEM/F12 and bacterial treatments were refreshed, and the TEER values were measured twice within each well, every 24 hours. Experiments were comprised of two biological replicates of each bacterial strain. Statistical differences in TEER were determined using the GraphPad Prism 9 software.

***Immunofluorescence (IF) staining and microscopy***. The ability of bacteria to prevent IFNγ-mediated loss of Zonula Occludens-1 (ZO-1) was determined by immunofluorescence staining and confocal microscopy. Briefly, T84 cells were cultured as previously outlined and seeded onto microscopy slides in 24-well plates at a density of 150,000 cells per well in 800 µL of DMEM/F12. Cells were grown for two weeks, and media refreshed every second day. The cells were pre-treated with 3 kDa-filtered medium extract (e.g. TY, PYG) or 3 kDa-filtered bacterial extract diluted to 1x in cell culture media and incubated at 37°C with 5% CO2. After pre-treatment for 1 h, 100 ng/mL IFNγ was added as appropriate and cells were incubated for a further 48 h. Cells were then fixed with 4% paraformaldehyde for 15 minutes, washed three times with PBS and permeabilised for 10 min with PBS supplemented with 0.1% v/v Triton X-100 and 5% w/v BSA. Cells were blocked in PBS supplemented with 5% w/v BSA for 1 h and the incubated with primary rabbit anti-human ZO-1 antibodies (5 µg/mL, Invitrogen) in PBS supplemented with 5% w/v BSA. The cells were then treated with secondary Alexa Fluor® 488-conjugated goat anti-rabbit IgG H&L antibodies (1:2000, ThermoFisher Scientific) and DAPI (1:1000, ThermoFisher Scientific) in PBS supplemented with 5% w/v BSA for 45 min. Coverslips were washed three times with PBS and once with ultrapure water before mounting on microscopy slides using ProLong Gold Antifade Mountant (Invitrogen). Image acquisition was performed using an inverted and fully motorised Nikon/Spectral Spinning Disc Confocal microscope (X-1 Yokogawa spinning disc with Borealis modification) with a 40x 1.30 Plan Fluor oil immersion objective lens (Nikon). Images were acquired using a coupled device (CCD) camera (Andow Clara) and the Nikon elements imaging software (Nikon, Version 4.40). For each condition, four separate images were captured of two biological replicates. Maximum projection images were assembled, and area of ZO-1 expression quantified and calculated for each image using Fiji ImageJ (Version 1.53t) and normalised to unstimulated control cells. Statistical analysis of differences in signal intensity were performed using the GraphPad Prism 9 software.

***Cell migration analysis.*** The Transwell® migration assays and IncuCyte® wound healing assays were used to assess the motility of human HCT116 gut epithelial cells during exposure to bacterial culture supernatant extracts. HCT116 cells were maintained in McCoys 5a medium supplemented with 10% FBS and 1% Pen/Strep.

To assess cell migration via the Transwell® assay, $3.5x10^4$ HCT116 cells were seeded in the apical compartment (100 µL volume) of a 6.5 mm insert with a TC-treated polycarbonate membrane in 24-well plates (8 µm pore size, Corning Costar). Then 600 µL of HCT116 culture medium was added to the lower compartment. The cells were allowed to settle for 24 hours. The following day, the cells were washed with Dulbecco's phosphate-buffered saline (DPBS), and then 100 µL of fresh culture medium with reduced serum content (0.5% FBS) was added to the top compartment and 600 µL of 0.5% FBS culture medium was added to lower compartment. Then, 0.5-2x concentrated extract from bacterial lead cultures were added to the lower compartment. After 16 hours, the HCT116 cells were washed with DPBS and the cells attached to the top of the membrane were carefully removed with a cotton tip. The migrated cells on the bottom of the membrane were then fixed in 70% ethanol for 10 minutes followed by staining in 0.25% crystal violet for 5 minutes. The Transwell® inserts were washed with water, dried and the membrane mounted with 50% glycerol in water on glass slides. The slides were imaged immediately and for each replicate two representative images of the membrane were taken at 10x magnification. The number of migrated cells was automatically counted using ImageJ and the average cell number displayed. The extent of cell migration was expressed as the average number of migrated cells in two microscopic fields per well from three biological and three technical replicates. Statistical differences in cell migration were determined using the GraphPad Prism 9 software.

For the IncuCyte® scratch wound assay, $3.5x10^4$ HCT116 cells were plated on poly-L-ornithine-coated IncuCyte® ImageLock 96-well plates (Essen BioScience). After 24 hours, the IncuCyte® WoundMaker tool was used to induce a homogeneous scratch wound in the nearly confluent cell monolayer. The cells were washed twice with DPBS and treated with 0.3x concentrated bacterial supernatant extracts diluted in 0.5% FBS McCoys 5a medium (200 µL total volume). Similarly prepared sterile bacterial medium extract served as negative control. Immediately after adding the stimulants, the plate was transferred to the IncuCyte® system and cell migration was monitored by imaging each well every two hours over the course of 72 hours. Data analysis was performed using the integrated analysis software. Statistical differences in cell migration were determined using the GraphPad Prism 9 software.

***Phage testing.*** Phage and prophage testing of the lead strains was performed by Phage Consultants (Gdansk, Poland). Briefly, a virulent phage test was performed by inoculating 250 µL of the sample into 25 mL of medium and incubating at 37ºC for 24 hours. Then, 250 µL of the culture was transferred to fresh medium and incubating at 37ºC for 24 hours. Both cultures were centrifuged at 4ºC for 15 minutes at 8000 rpm in a Sigma 3-18K centrifuge and then 10 mL of the collected supernatant was filtered using a 0.22 µm syringe filter. Next, 10 µL of each sample was prepared for electron microscopy by adding 500 U of viscolase. A control sample was prepared by adding $10^9$ crude T4 lysate to the sample. For the mitomycin C induction test, 250 µL of the sample was inoculated into 25 mL of medium and incubated at 37ºC for 3 hours after which 1 µg/ml of mitomycin C was added. After 24 hours, the culture was centrifuged at 4ºC for 15 minutes at 8000 rpm in a Sigma 3-18K centrifuge. Then, 10 mL of the collected supernatant was filtered using 0.22 µm syringe filter. Next, 10 µL of the sample was prepared for electron microscopy by adding 500 U of viscolase.

Electron microscopy was performed using a Tecnai Spirit BioTWIN electron microscope with a digital camera. The samples were prepared for negative staining by placement on C400Cu100 grids (400 Mesh, copper and carbon layer coating) manufactured by EM Resolutions. Samples were negative stained with 2% uranyl acetate in water at room temperature for 15 seconds. In the absence of visible phage particles, only general view images were captured.

***Antibiotic susceptibility testing.*** Antibiotic susceptibility testing was performed by Mater Pathology (Brisbane, Australia). Briefly, a ~1 McFarland suspension of each strain was prepared in Schaedler broth. The suspensions were plated on anaerobic blood agar except for MH35 which was plated on TY based medium (TY medium modified to include menadione (100 µg/mL) and lack haemin). Susceptibility testing to amoxicillin, chloramphenicol, clindamycin and vancomycin was performed using Etests and results were interpreted using EUCAST's species agnostic anaerobe breakpoints. The minimum inhibitory concentration for other antibiotics was determined using the Etest instructions for use.

***Statistical and data analyses.*** Data are presented as mean with standard deviation (SD). Statistical analysis for each figure were preformed using GraphPad Prism V9 software. All statistical tests were two-sided unless stated otherwise. Details of the statistical tests and methodology for quantification are provided in the figure legends.

| Component | *H. mulieris* custom medium (g/L) | *H. fusiformis* custom medium (g/L) | *V. sanitatis* custom medium (g/L) |
|---|---|---|---|
| Trehalose | 1 g | | |
| Xylose | | 1 g | |
| Glycine | | 0.2 g | |
| Tryptophan | 0.08 g | 0.09 g | 0.1 g |
| Methionine | 0.2 g | 0.2 g | 0.2 g |
| Leucine | | 0.2 g | 0.2 g |
| Proline | | 0.2 g | |
| Serine | | 0.2 g | |
| Threonine | | 0.2 g | |
| Phenylalanine | 0.2 g | 0.2 g | 0.2g |
| Tyrosine | 0.2 g | 0.2 g | 0.2 g |
| Histidine | | 0.125 g | 0.2 g |
| Lysine hydrochloride | | 0.725 g | 0.725 g |
| Alanine | 0.4 g | | 0.2 g |
| Arginine | | | 0.2 g |
| Isoleucine | | | 0.2 g |
| Valine | | | 0.2 g |
| *SCFA mix | | 3.1 mL | 3.1 mL |
| Butyric acid | 400 µL | | |
| Sodium azide (10% w/v) | | | 2.5 mL |
| Salts solution 2 [24] | 75 mL | 38 mL | 38 mL |
| Salts solution 3 [24] | 75 mL | 38 mL | 38 mL |
| Sodium bicarbonate | 8 g | 8 g | 8 g |
| Resazurin (0.1 w/v) | 1 mL | 1 mL | 1 mL |
| L-Cysteine | 1 g | 1 g | 1 g |
| *Vitamin solution (1000x) | 1 mL | 1 mL | 1 mL |

**Supplementary Table S1.** Custom GDI media designed for *H. mulieris*, *H. fusiformis* and *V. sanitatis*. GDI media wss comprised of core components (salt solutions 2 &3, sodium bicarbonate, resazurin, L-cysteine and vitamin solution). The core GDI medium wass supplemented with other components based on the GDI predictions. Sodium azide was added to suppress growth of Gram-negative bacteria.

*SCFA (short chain fatty acid) mix: Acetic acid 17mL/L, Propionic acid 6mL/L, Butyric acid 4ml/L, Isobutyric acid 1mL/L, Valeric acid 1mL/L, Isovaleric acid 1mL/L and 2-methylbutyric acid 1mL/L made up in 1L $H_2O$

*Vitamin solution (1000X): Pyridoxine HCL 100 µg/L, L-Ascorbic acid 100 µg/L, Calcium pantothenate 100 µg/L, Nicotinamide 100 µg/L, Pyridoxal 100 µg/L, Thiamine – HCL 100 µg/L, Nicotinic acid 100 µg/L, D-Biotin 100 µg/L, P-aminobenzoic acid 100 µg/L, Vitamin B12 100 µg/L, Riboflavin 50 µg/L, Folic acid 100 µg/L.

| Feature | Scoring |
|---|---|
| Thickening of the colon wall | |
| Transparent | 0 |
| Moderate | 1 |
| Marked | 2 |
| Intransparent | 3 |
| Changes in vascular pattern | |
| Normal | 0 |
| Moderate | 1 |
| Marked | 2 |
| Bleeding | 3 |
| Granularity of the mucosal surface | |
| None | 0 |
| Moderate | 1 |
| Marked | 2 |
| Extreme | 3 |
| Stool consistency/mucus secretion | |
| Normal and solid | 0 |
| Still shaped, mild mucus | 1 |
| Unshaped, mucus | 2 |
| Spread | 3 |
| Extent of involved area | |
| 0-5% (none) | 0 |
| 5-20% (patchy) | 1 |
| 20-50% (moderate) | 2 |
| >50% (predominant) | 3 |

**Supplementary Table S2.** Scoring of gut barrier dysfunction by colonoscopy[38,39].

| | |
|---|---|
| **Inflammation score (Scored 0-4)** | |
| 0 | No evidence of inflammation |
| 1 | Low level of inflammation with scattered infiltrating mononuclear cells (1-2 foci only) |
| 2 | Moderate inflammation with multiple foci |
| 3 | High level of inflammation with increased vascular density and marked wall thickening |
| 4 | Maximal severity of inflammation with transmural leukocyte infiltration and loss of goblet cells |
| **Injury score (Scored 0-3)** | |
| 0 | No epithelial injury |
| 1 | Occasional epithelial lesion |
| 2 | 1-2 foci of ulceration |
| 3 | Extensive ulceration |
| **Colitis activity (Composite score (/17) based on measures listed below)** | |
| Hypervascularisation | 0-3 based on severity |
| Presence of mononuclear cells | 0-3 based on severity |
| Epithelial hyperplasia | 0-3 based on severity |
| Epithelial injury | 0-3 based on severity |
| Presence of neutrophils | 0-3 based on severity |
| Lymphoid aggregates | Scored 0-2, where 0 = 0, 1 = ≤2, and 2 = ≥2 |

**Supplementary Table S3.** Scoring of histological colitis[38,39]. The epithelial injury score is a composite score of epithelial hyperplasia and epithelial injury. The inflammation score is a composite of presence of mononuclear cells, neutrophils and lymphoid aggregates.

| Feature | Grade | Description |
|---|---|---|
| **Inflammatory infiltrate** | 0 | Absent |
| | 1 | Scattered PMNs or MNs in lamina propria (LP) involving <5 intercryptal spaces contiguously involved in an area |
| | 2 | Increased PMNs or MNs in LP and/or submucosa involving >5 intercryptal spaces contiguously involved in an area and ± crypt abscesses and areas of crypt/villous loss < 5 crypt widths |
| | 3 | Extensive transmural inflammatory infiltrate and areas of crypt/villous loss > 5 crypt widths and/or confluent areas of crypt abscesses and/or mucosal erosion/ulceration |
| **Granulomata** | 0 | Absent |
| | 1 | Aggregation of cells not definitely a granuloma |
| | 2 | A definite granuloma formation present |
| | 3 | Multiple granulomas present |
| **Villous Distortion** | 0 | Absent |
| | 1 | Villus height increased or decreased by <1/3 of normal |
| | 2 | Villus height increased or decreased by 1/3-2/3 of normal |
| | 3 | Villus height increased or decreased by >2/3 of normal |
| **Cross sectional area** | **Score** | |
| <1% | 0.5 | |
| 1-25% | 1 | |
| 26-50% | 2 | |
| 51-75% | 3 | |
| >75% | 4 | |

**Supplementary Table S4.** Histological scoring system for ileum of SKG mice[40]. For total histology score add scores from sections 1, 2 and 3 and multiply by cross sectional area involved.

| Class | Biogenic amine | Pathway definitions | Enzyme descriptions |
|---|---|---|---|
| **Aromatic/Heterocyclic amines** | Histidine | **4.1.1.22** | Histidine decarboxylase |
| | Tyramine | **4.1.1.25**/ 4.1.1.28 | Tyrosine decarboxylase |
| | Tryptamine | **4.1.1.28**/ 4.1.1.105 | Tryptophan decarboxylase |
| | Phenylethylamine | **4.1.1.28**/ 4.1.1.53 | Phenylalanine decarboxylase |
| | Serotonin | **4.1.1.28** | 5-hydroxy-L-tryptophan decarboxylase |
| **Aliphatic amines** | Agmatine | **4.1.1.19** | Arginine decarboxylase |
| | Cadaverine | **4.1.1.18** | Lysine decarboxylase |
| | Putrecine_1 | 4.1.1.19 + 3.5.3.12 + **3.5.1.53** | Arginine decarboxylase, Agmatine deiminase, N-carbamoylputreciscine aminohydrolase |
| | Putrecine_2 | 4.1.1.19 + **3.5.3.11** | Arginine decarboxylase, Agmatinase |
| | Putrecine_3 | 3.5.3.1 + **4.1.1.17** | Arginase, Ornithine decarboxylase |
| | Spermidine_1 | 4.1.1.50 + **2.5.1.16** | S-adenosylmethionine decarboxylase, Spermidine synthase |
| | Spermidine_2 | 2.7.4.2 + 1.2.1.11 + 1.5.1.43 + **4.1.1.96** | Aspartate kinase Aspartate-beta-semialdehyde dehydrogenase Carboxyspermidine dehydrogenases Carboxyspermidine decarboxylase |
| | Spermidine_3 | 4.1.1.9 + 4.1.1.50 + 2.5.1.104 + **3.5.3.24** | Pyruvoyl-dependent arginine decarboxylase S-adenosylmethionine decarboxylase Agmatine aminopropyltransferase $N^1$-aminopropylagmatine ureohydrolase |
| | Spermine | 4.1.1.50 + **2.5.1.22** | S-adenosylmethionine decarboxylase Spermine synthase |

**Supplementary Table S5.** Pathway definitions for key biogenic amines. + indicates that an enzyme is required in addition to the following enzymes, while an '/' indicates the enzyme is interchangeable with the following enzyme.

| | | *A. shahii* | *A. communis* | *M. faecis* | | *H. mulieris* | *G. formicilis* | *H. fusiformis* | | *V. sanitatis* |
|---|---|---|---|---|---|---|---|---|---|---|
| | | MH21-1 | MH22-1 | MH23-1 | MH23-3 | MH27-1 | MH32-1 | MH35-1 | MH35-2 | MH37-1 |
| Antibiotic resistance | Tetracycline (is / iv) | S/0.25 | R/64 | R/16 | S/0.25 | R/0.016 | R/0.016 | R/1 | R/1 | R/- |
| | Chloramphenicol (is / iv) | R/R | S/S | S/S | S/S | S/S | S/S | S/S | S/S | S/n.t. |
| | Vancomycin (is / iv) | S/n.t.* | S/n.t.* | S/S | !/S | S/S | S/S | S/S | S/S | R/n.t. |
| | Bacitracin (is / iv) | S/n.t. | S/n.t. | S/n.t. | S/n.t. | S/n.t. | S/n.t. | R/n.t. | S/n.t. | S/n.t. |
| | Amoxicillin (is / iv) | S/S | R/R | !/S | !/S | S/S | S/S | S/0.125 | S/0.016 | S/n.t. |
| | Clindamycin (is / iv) | S/S | S/S | R/S | S/S | R/S | S/S | S/S | S/S | S/n.t. |
| | Metronidazole (is / iv) | S/S | S/S | S/S | S/S | S/S | S/S | S/S | S/S | S/n.t. |
| Virulence | All (is) | n.d. | n.d. | n.d. | n.d. | n.d. | n.d. | n.d. | n.d. | n.d. |
| Metabolites | Harmful biogenic amines (is) | n.d. | n.d. | n.d. | n.d. | n.d. | n.d. | n.d. | n.d. | n.d. |
| Mobile elements | Plasmid (is) | n.d. | n.d. | 1 | n.d. | n.d. | 1 | n.d. | 1 | 1 |
| | Phage (iv) | n.d. | n.d. | n.d. | n.d. | n.d. | + | n.d. | n.d. | n.d. |
| | Mobile elements (is) | 2 | n.d. | n.d. | 1 | 1 | n.d. | n.d. | n.d. | n.d. |

**Supplementary Table S6.** Summary of *in silico* and *in vitro* safety assessments for the IBD therapeutic leads. Leads were also assessed for the presence of phage, which can impact manufacturing. is = *in silico*, iv = *in vitro*, S = Sensitive, R = Resistant, ! = Regulatory genes detected. n.t. = not tested, - = not detected, * = vancomycin does not affect gram negative strains.

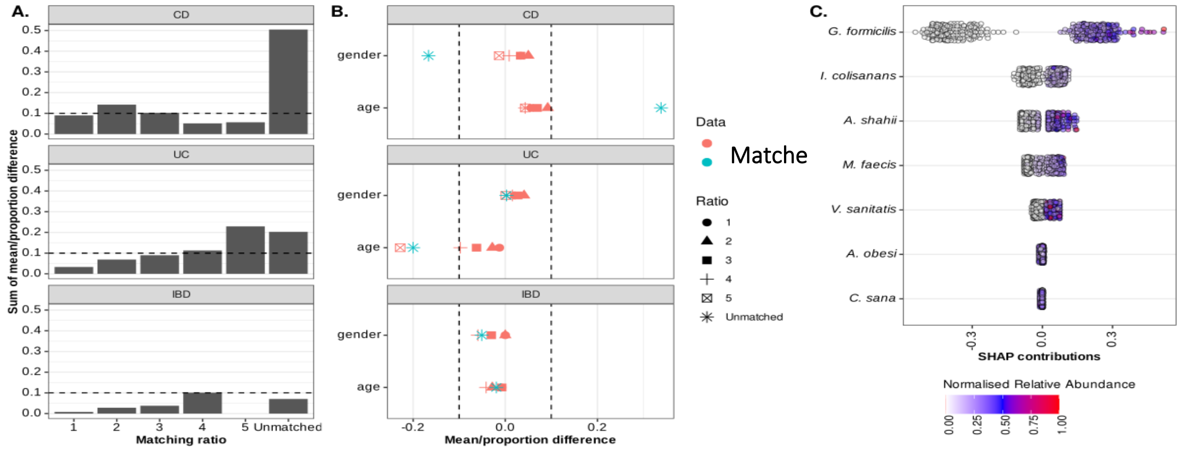


**Supplementary Figure 1.** (A) Matching of study groups from the Australian cohort by age, gender and BMI. Healthy controls were separately matched to IBD, CD and UC cohorts using propensity score matching. Shown is the sum of standard mean difference (SMD) for age, and difference in proportion for gender, before (Unmatched) and after matching for matching ratios (1:1 – 1:5). Highest ratios at which the sum of SMD and difference in proportion was <0.1 are highlighted by red stars. (B) Love plot showing the individual standard mean difference for gender and age, and difference in proportion for BMI at each of the assessed matching ratios. Shown are the SMD and difference in proportion before matching (unadjusted) and after matching (adjusted) at different matching ratios. (C) Lead species SHAP contributions for the gradient boosting machine learning model trained to discriminate controls from individuals with IBD in a combined Australian and US cohort (Figure 2A).

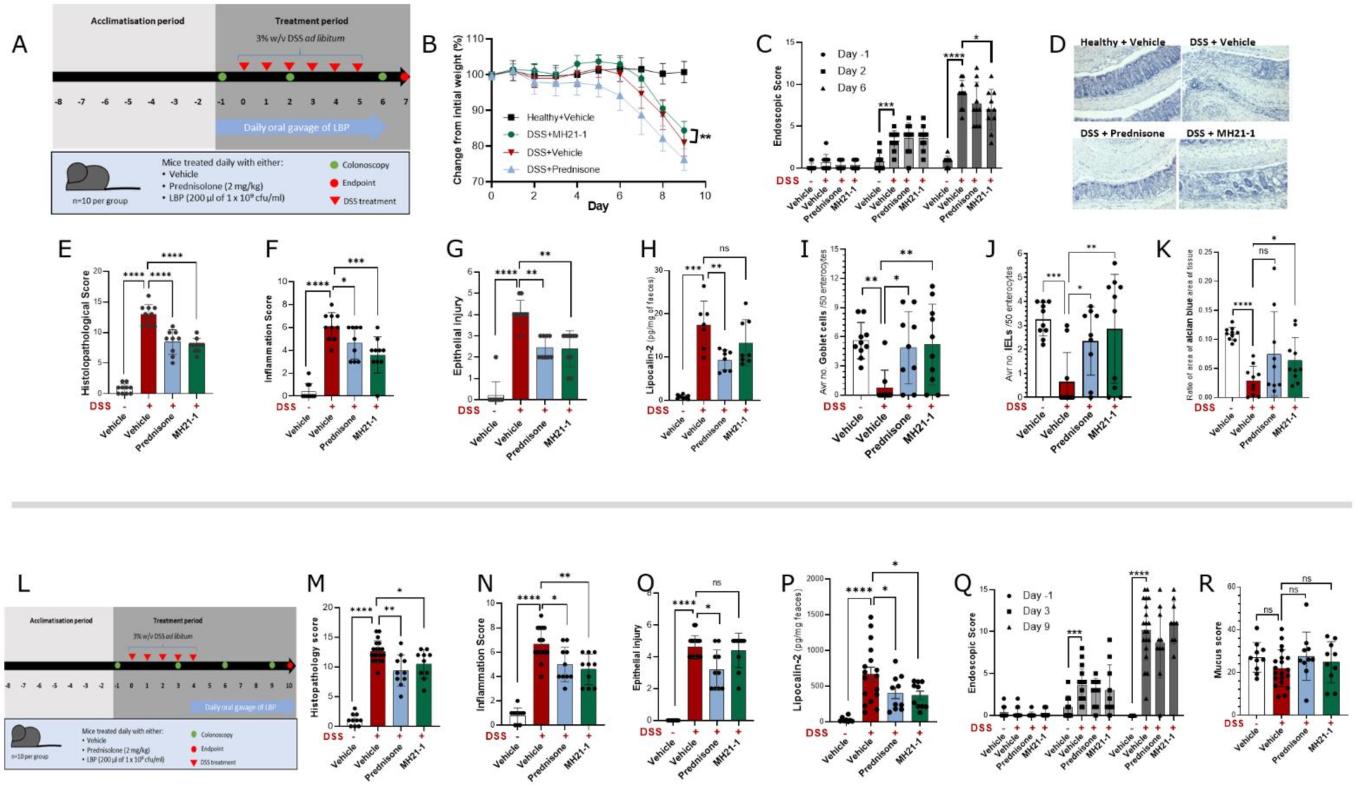

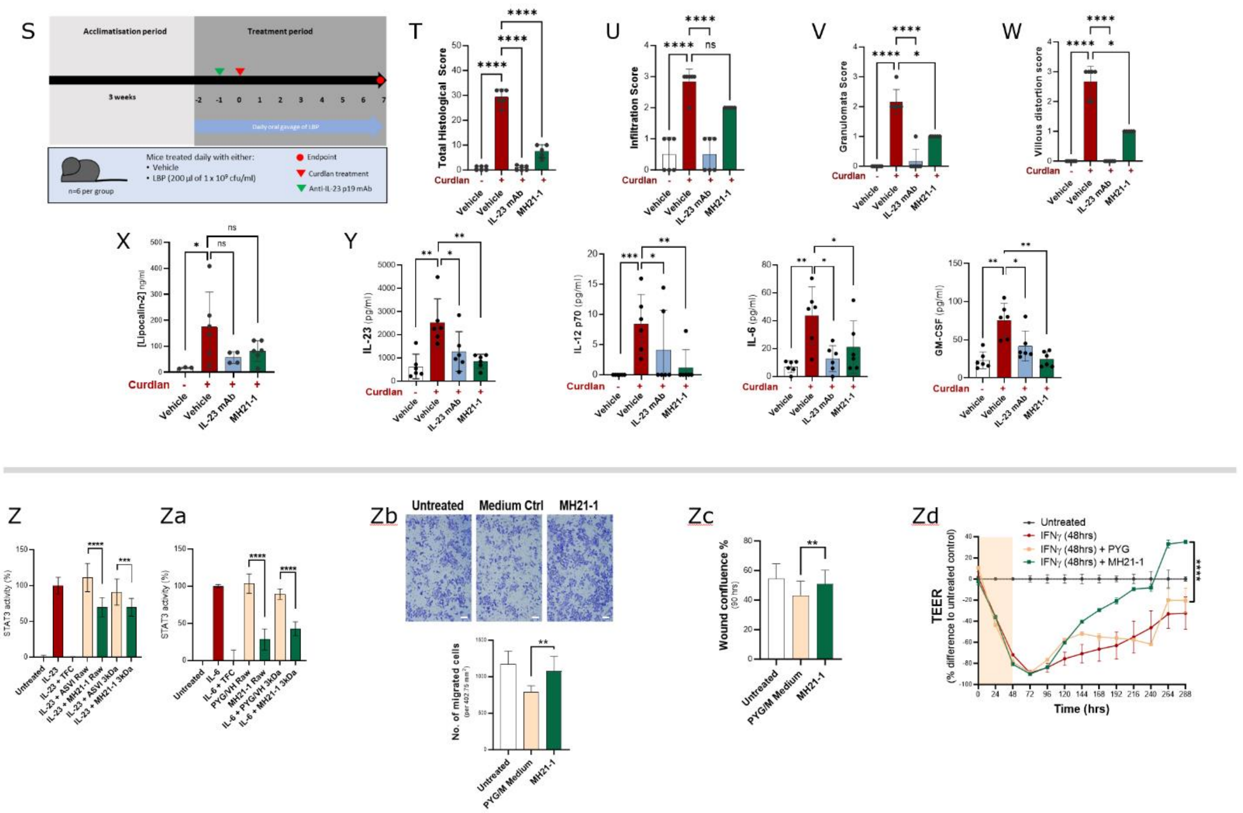


**Supplementary Figure 2.**

Assessment of therapeutic activity of *A. shahii* MH21-1. Live cultures of MH21-1 were assessed in a prophylactic DSS mouse model (A). MH21-1 treatment reduced weight loss in mice receiving DSS (B$^{\#}$). MH21-1 improved the endoscopic score compared to vehicle in DSS receiving mice on day 6 of treatment (C, $^{\$}$day -1, $^{\%}$day 2, 6). MH21-1 improved DSS induced colitis as illustrated by representative gut histology images (D) and ameliorated increases in histopathological score (E$^{\&}$), inflammation (F$^{\&}$) and epithelial injury (G$^{\$}$) compared to vehicle control. MH21-1 did not significantly reduce the lipocalin-2 concentrations in faeces of DSS-treated mice compared to mice treated with DSS and vehicle (H$^{\&}$). DSS treatment resulted in a significant decrease in goblet cells, intraepithelial lymphocytes and mucus production that was ameliorated by treatment with MH21-1 (I$^{\$}$, J$^{\$}$, K$^{\&}$). Study design of the therapeutic DSS mouse model (L). In this model, MH21-1 ameliorated increases in histopathological score (M$^{\&}$), inflammation (N$^{\$}$) but not epithelial injury (O$^{\$}$) compared to the vehicle control. MH21-1 ameliorated the increases in faecal lipocalin-2 compared to mice treated with DSS and vehicle (P$^{\&}$). MH21-1 treatment did not result in significant improvements in the endoscopic or mucus score compared to mice receiving DSS and vehicle (Q$^{\$}$, R$^{\%}$). MH21-1 was assessed in SKG mouse model of ileitis (S). Curdlan treatment resulted in an increase in the histopathological and infiltration score that was ameliorated by treatment with MH21-1 (T$^{\&}$, U$^{\$}$). MH21-1 did not significantly improve the increased granulomata (V$^{\$}$) and villous distortion score (W$^{\$}$), nor the higher lipocalin-2 levels (X$^{\&}$) resulting from curdlan treatment. Curdlan treatment resulted in an increase in IL-23, IL-12p70, IL-6 and GM-CSF that was ameliorated by MH21-1 (Y; $^{\&}$for IL-23, IL-6 and GM-CSF; $^{\$}$ for IL-12). MH21-1 culture supernatant suppressed IL-23-mediated STAT3 activation in HEKBlue IL-23 reporter cells (Z$^{\wedge}$). MH21-1 culture supernatant suppressed IL-6-mediated STAT3 activation in HEKBlue IL-6 reporter cells (Za$^{\wedge}$). MH21-1 significantly increased the migration of HCT116 cells compared to the medium control in a Transwell migration assay (Zb$^{\wedge}$; scale bar 100 μm). MH21-1 supernatant extract promoted faster wound closure of HCT116 cells compared to the controls in an IncuCyte scratch wound assay (Zc$^{\wedge}$ 90 hrs post scratch). MH21-1 promoted faster recovery from IFNγ-mediated barrier integrity loss as indicated by TEER across confluent monolayers of T84 gut epithelial cells compared to the medium control (Zd; t-test at endpoint 288 hrs). For all data, ns: not significant; *, $p < 0.05$; **, $p < 0.01$; ***, $p < 0.001$; ****, $p < 0.0001$. All data presented as mean and standard deviation. [#]Two-way ANOVA with Fisher's test for multiple comparison. [$]Kruskal-Wallis test with uncorrected Dunn's for multiple comparisons. [&]Uncorrected Brown-Forsythe and Welch ANOVA test with multiple comparisons. [%]One-way ANOVA with uncorrected Fisher's LSD test for multiple comparison. [^]T-test medium control vs lead.

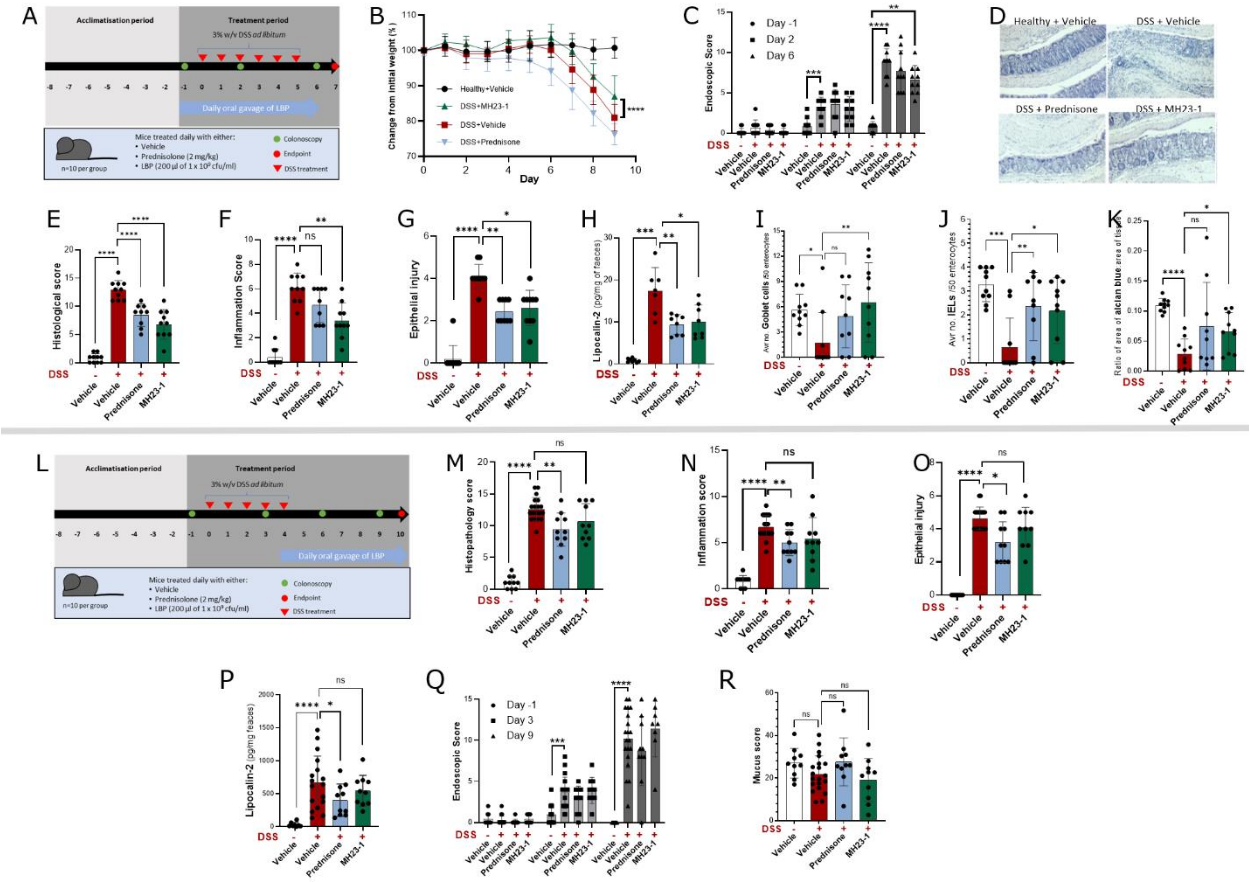

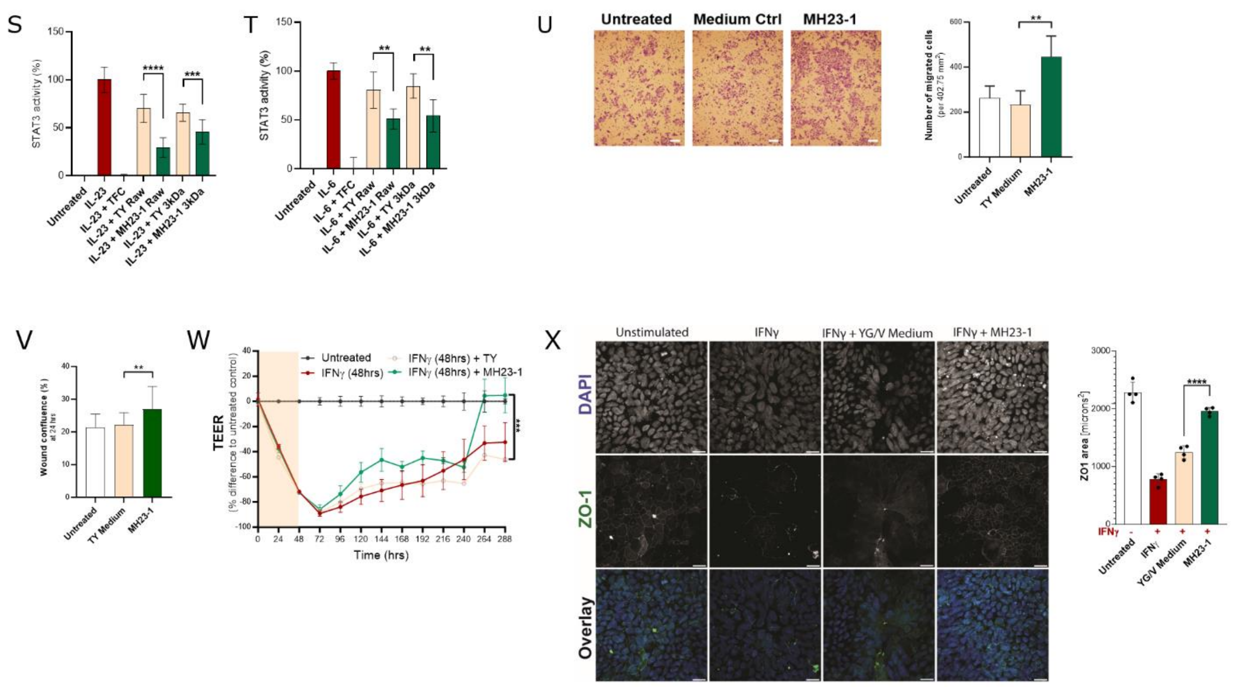


**Supplementary Figure 3.**

Assessment of therapeutic activity of *M. faecis* MH23-1. Live cultures of MH23-1 were assessed in a prophylactic DSS mouse model (A). MH23-1 treatment relieved weight loss in mice receiving DSS (B[#]). MH23-1 improved the endoscopic score compared to vehicle in DSS receiving mice on day 6 of treatment (C, [$]day -1, [%]day 2, 6). MH23-1 improved DSS induced colitis as illustrated by representative gut histology images (D) and ameliorated increases in histopathological score (E[&]), inflammation (F[$]) and epithelial injury (G[$]) compared to vehicle control. MH23-1 reduced the faecal lipocalin-2 concentrations in mice treated with DSS compared to the control, DSS plus vehicle (H[&]). DSS treatment resulted in a significant decrease in goblet cells, intraepithelial lymphocytes and mucus production that was ameliorated by MH23-1 (I[$], J[$], K[&]). Study design of therapeutic DSS mouse model (L). In this model, MH23-1 did not ameliorate increases in histopathological score (M[&]), inflammation (N[&]) or epithelial injury (O[$]) compared to the vehicle control. MH23-1 did not affect faecal lipocalin-2 concentrations (P[&]). MH23-1 treatment did not result in improvements in the endoscopic or mucus score compared to mice receiving DSS and vehicle (Q[$], R[%]). MH23-1 culture supernatant suppressed IL-23-mediated STAT3 activation in HEKBlue IL-23 reporter cells (S[^]). MH23-1 culture supernatant suppressed IL-6-mediated STAT3 activation in HEKBlue IL-6 reporter cells (T[^]). MH23-1 significantly increased the migration of HCT116 cells compared to the medium control in a Transwell migration assay (U[^]; scale bar 100 μm). MH23-1 supernatant extract promoted faster wound closure of HCT116 cells compared to the controls in an IncuCyte scratch wound assay (V[^] 24 hrs post scratch). MH23-1 promoted faster recovery from IFNγ-mediated barrier integrity loss as indicated by TEER across confluent monolayers of T84 gut epithelial cells compared to the medium control (W[^] at endpoint 288 hrs). MH27-1 ameliorated IFNγ-induced reductions in ZO-1 expression in T84 cells compared to the medium control (X[^]; scale bar 100 μm) For all data, ns: not significant; *, $p < 0.05$; **, $p < 0.01$; ***, $p < 0.001$; ****, $p < 0.0001$. All data presented as mean and standard deviation. [#]Two-way ANOVA with Fisher's test for multiple comparison. [$]Kruskal-Wallis test with uncorrected Dunn's for multiple comparisons. [&]Uncorrected Brown-Forsythe and Welch ANOVA test with multiple comparisons. [%]One-way ANOVA with uncorrected Fisher's LSD test for multiple comparison. [^]T-test of medium control vs lead.

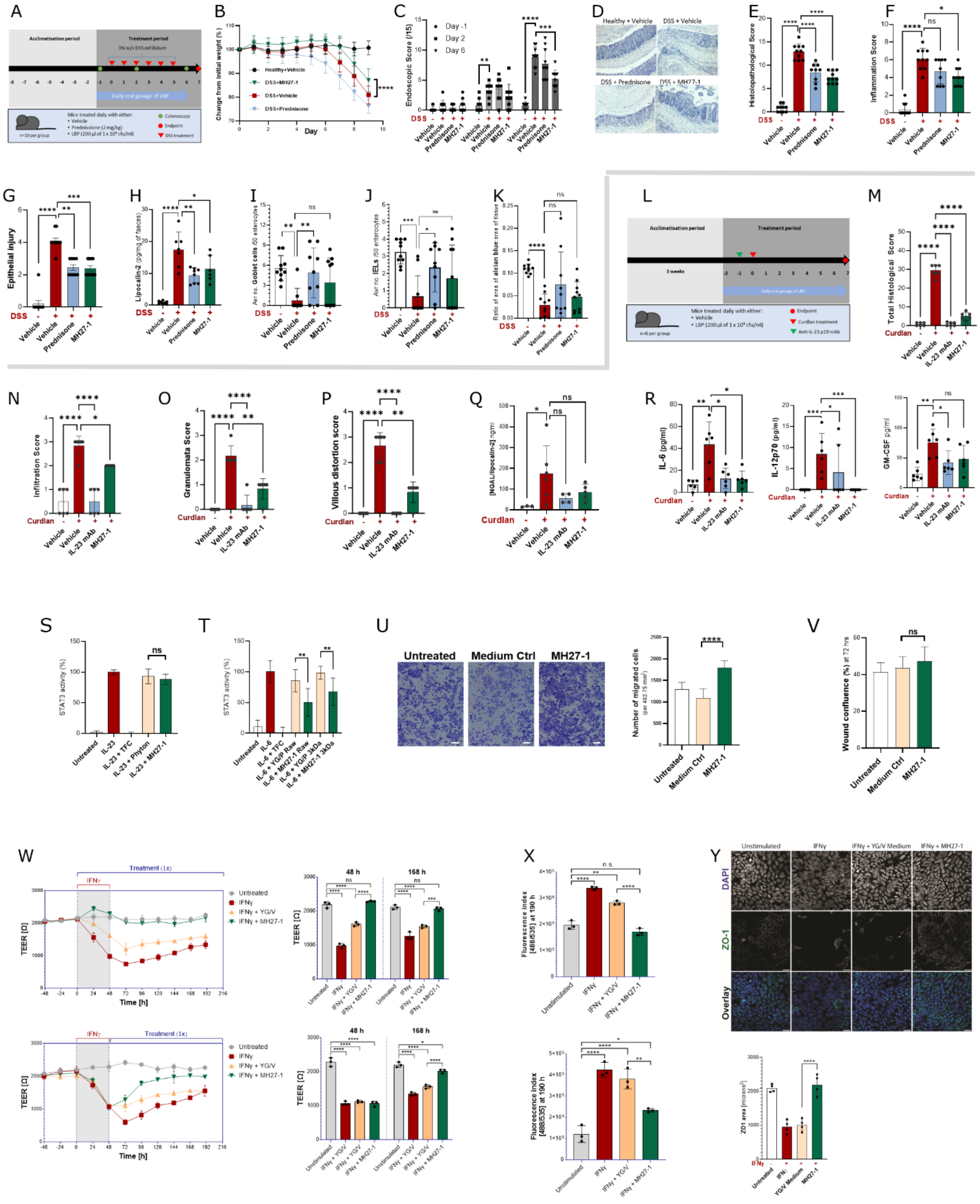


**Supplementary Figure 4.**

Assessment of therapeutic activity of *H. mulieris* MH27-1. Live cultures were assessed in a prophylactic DSS mouse model (A). MH27-1 treatment relieved weight loss in mice receiving DSS (B$^{\#}$). MH27-1 improved the endoscopic score compared to vehicle in DSS receiving mice on day 6 of treatment (C, $^{\$}$day

-1, [%]day 2, 6). MH27-1 improved DSS induced colitis as illustrated by representative gut histology images (D) and ameliorated increases in histopathological score (E[&]), inflammation (F[$]) and epithelial injury (G[$]) compared to vehicle control. Compared to DSS and vehicle, faecal lipocalin-2 levels were significantly lower in mice treated with DSS and MH27-1 (H[&]). DSS treatment resulted in a significant decrease in goblet cells, intraepithelial lymphocytes and mucus production, and these readouts were not affected by treatment with MH27-1 (I[$], J[$], K[&]). MH27-1 was assessed in SKG mouse model of ileitis (L). Curdlan treatment resulted in an increase in the histopathological and infiltration score that was ameliorated by treatment with MH27-1 (M[&], N[$]). MH27-1 did not significantly improve the increased granulomata (O[$]) and villous distortion score (P[$]), nor the higher lipocalin-2 levels (Q[&]), resulting from curdlan treatment. Curdlan treatment resulted in an increase in IL-12p70, IL-6 and GM-CSF that were significantly ameliorated by MH27-1 for IL-12 and IL-6, compared to the DSS + vehicle group (R; [&] for IL-6 and GM-CSF; [$] for IL-12). Culture supernatant of MH27-1 did not impact IL-23-mediated STAT3 signalling (S[^]). IL-6-mediated STAT3 signalling was inhibited in HEKBlue IL-6 reporter cells treated with culture supernatant of MH27-1 compared to the medium control (T[^]). Compared to the medium extract control, culture supernatant extract from MH27-1 promoted the migration of HCT116 colon cancer cells in the Transwell migration assay (U[^]; scale bar 100 μm). MH27-1 extract did not promote accelerated wound closure in the IncuCyte scratch wound assay compared to cells treated with medium extract (V[^]). As indicated by measuring TEER across confluent monolayers of T84 gut epithelial cells, MH27-1 prevented IFNγ-mediated barrier integrity loss in a prophylactic model and promoted faster recovery from IFNγ insult in a therapeutic model, compared to the medium control (W[^]). MH27-1 ameliorates IFNγ-mediated changes in cellular permeability as evidenced by movement of fluorescent labelled dextran across T84 cell monolayers (X[^]). MH27-1 ameliorated IFNγ-induced reductions in ZO-1 expression in T84 cells compared to the medium control (Y[^]; scale bar 100 μm). For all data, ns: not significant; *, $p < 0.05$; **, $p < 0.01$; ***, $p < 0.001$; ****, $p < 0.0001$. All data presented as mean and standard deviation. [#]Two-way ANOVA with Fisher's test for multiple comparison. [$]Kruskal-Wallis test with uncorrected Dunn's for multiple comparisons. [&]Uncorrected Brown-Forsythe and Welch ANOVA test with multiple comparisons. [%]One-way ANOVA with uncorrected Fisher's LSD test for multiple comparison. [^]T-test of medium control vs lead.

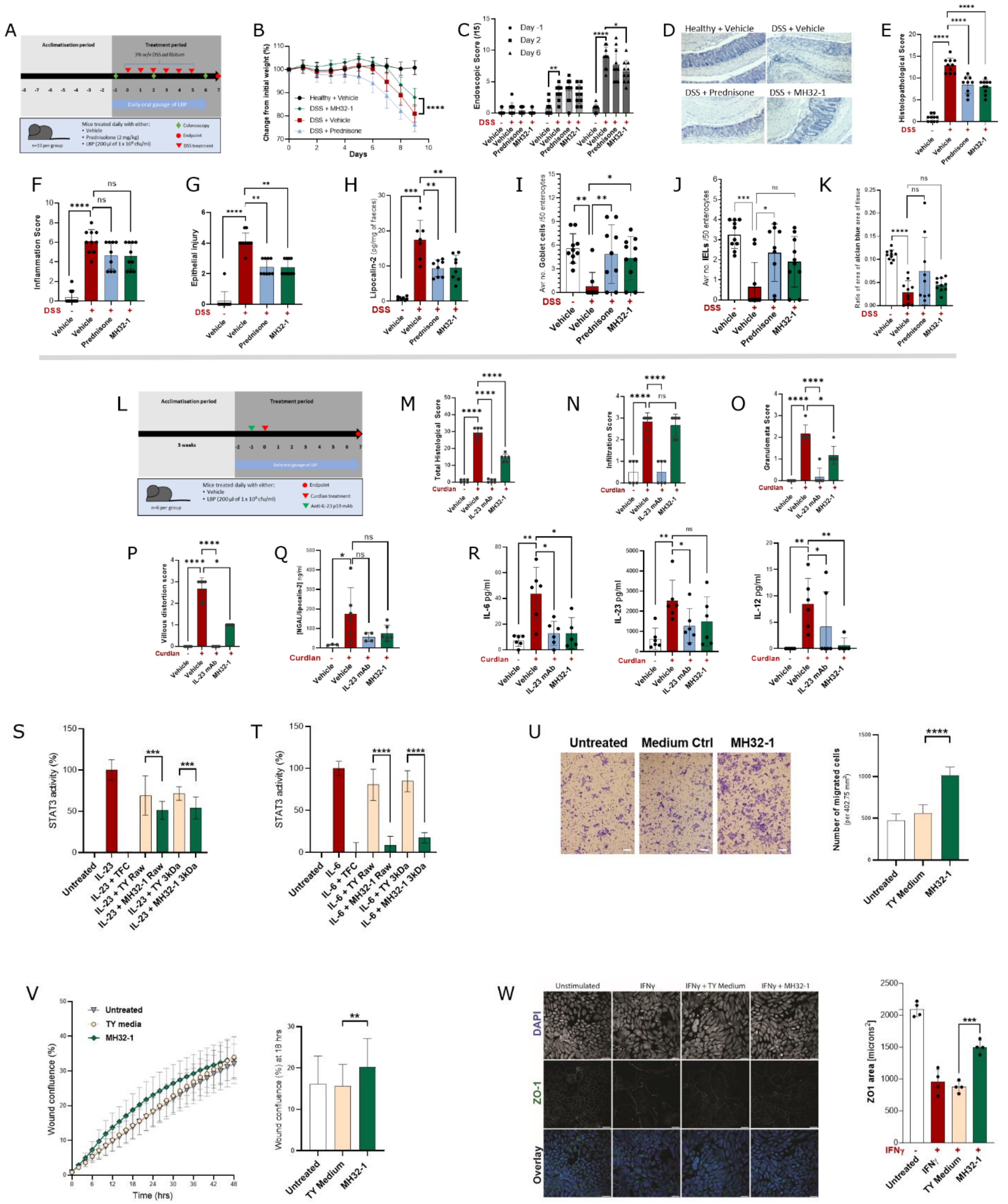


**Supplementary Figure 5.**

Assessment of therapeutic activity of *G. formicilis* MH32-1. Live cultures of MH32-1 were assessed in a prophylactic DSS mouse model (A). MH32-1 treatment relieved weight loss in mice receiving DSS compared to the DSS + vehicle group (B$^{\#}$). MH32-1 improved the endoscopic score compared to vehicle in DSS receiving mice on day 6 of treatment (C, $^{\$}$day -1, $^{\%}$day 2, 6). MH32-1 improved DSS induced colitis as illustrated by representative gut histology images (D) and ameliorated increases in histopathological score (E$^{\&}$), inflammation (F$^{\$}$) and epithelial injury (G$^{\$}$) compared to vehicle control, however, the change in inflammation score did not reach statistical significance. Compared to mice treated with DSS + vehicle, lipocalin-2 concentrations in faeces were significantly lower in mice treated with DSS and MH32-1 (H$^{\&}$). DSS treatment resulted in a significant decrease in goblet cells, intraepithelial lymphocytes and mucus production and the reduction in goblet cells was significantly ameliorated by concurrent treatment with MH32-1 (I$^{\$}$, J$^{\$}$, K$^{\&}$). MH32-1 was assessed in the SKG mouse model of ileitis (L). MH32-1 ameliorated the curdlan-mediated increase in the histopathological score but not the inflammation score when compared to the curdlan + vehicle group (M$^{\&}$, N$^{\$}$). MH32-1 did not significantly improve the increased granulomata (O$^{\$}$) and villous distortion score (P$^{\$}$), nor the higher lipocalin-2 levels (Q$^{\&}$) resulting from curdlan treatment. Curdlan treatment resulted in an increase in IL-6, IL-23 and IL-12 that was significantly ameliorated for IL-6 and IL-12 by treatment with MH32-1 when compared to the DSS + vehicle group (R; $^{\&}$ for IL-6 and IL-23; $^{\$}$ for IL-12). Treatment with culture supernatant of MH32-1 had a small but significant suppressive effect on IL-23-mediated STAT3 signalling in HEKBlue IL-23 reporter cells (S$^{\wedge}$). IL-6-induced STAT3 signalling was inhibited when HEKBlue IL-6 reporter cells were treated with culture supernatant of MH32-1 (T$^{\wedge}$). MH32-1 extract significantly increased the movement of HCT116 gut epithelial cells compared to medium extract control in the Transwell migration assay (U$^{\wedge}$; scale bar 100 µmHCT116 cells incubated in extract from MH32-1 showed significantly higher wound confluence, compared to cells treated with TY medium extract, at 18 hrs post scratch, in the IncuCyte scratch wound assay (V$^{\wedge}$). MH32-1 ameliorated IFNγ-induced reductions in ZO-1 protein expression in T84 cells compared to the medium control (W$^{\wedge}$; scale bar 100 µm). For all data, ns: not significant; *, $p < 0.05$; **, $p < 0.01$; ***, $p < 0.001$; ****, $p < 0.0001$. All data presented as mean and standard deviation. $^{\#}$Two-way ANOVA with Fisher's test for multiple comparison. $^{\$}$Kruskal-Wallis test with uncorrected Dunn's for multiple comparisons. $^{\&}$Uncorrected Brown-Forsythe and Welch ANOVA test with multiple comparisons. $^{\%}$One-way ANOVA with uncorrected Fisher's LSD test for multiple comparison. $^{\wedge}$T-test medium control vs lead.

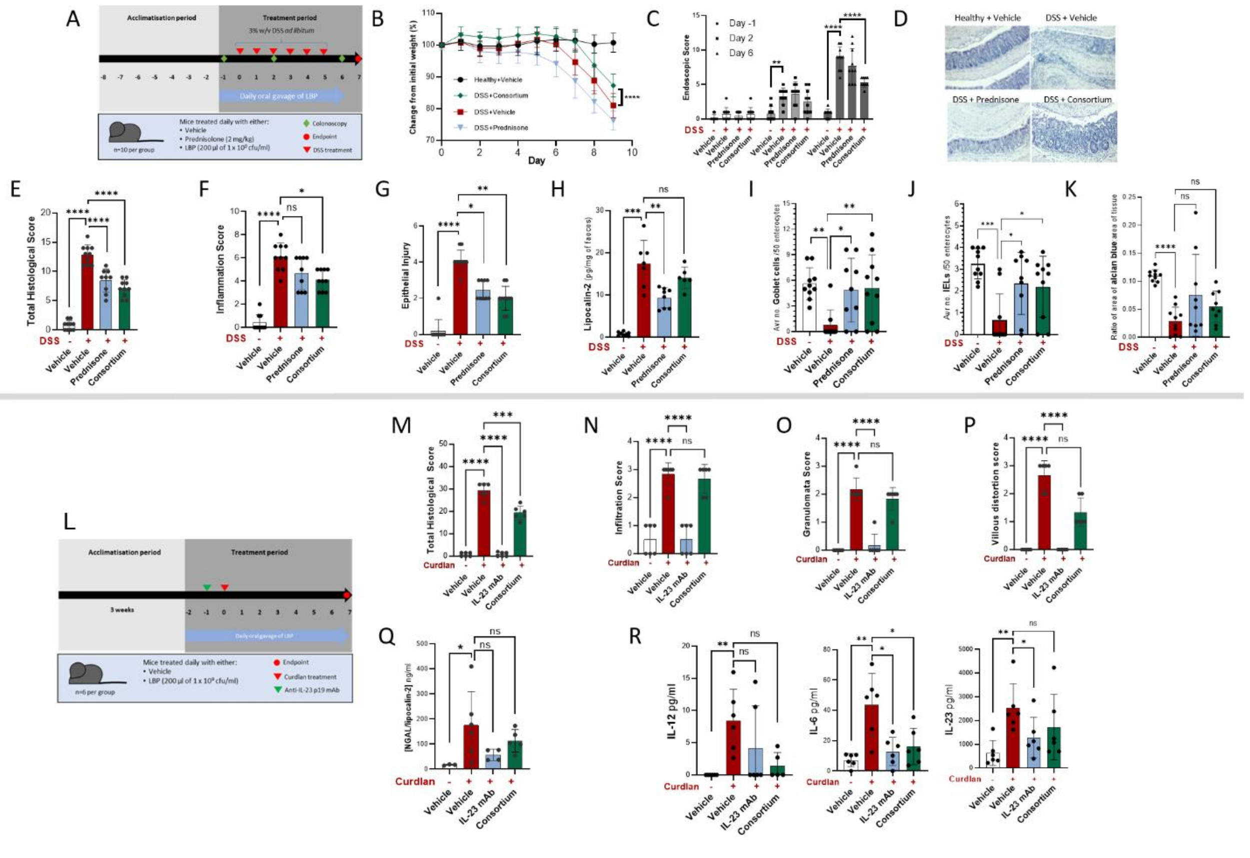

**Supplementary Figure 6.**

Assessment of therapeutic activity of the consortium MAP#4 of the four lead candidates *A. shahii* MH21-1, *M. faecis* MH23-1, *H. mulieris* MH27-1, and *G. formicilis* MH32-1. (A) Prophylactic DSS mouse model of murine colitis. Treatment with consortium MAP#4 relieved weight loss in mice receiving DSS compared to the DSS + vehicle group (B#). Treatment with MAP#4 improved the endoscopic score compared to vehicle in DSS receiving mice on day 6 of treatment (C, $day -1, %day 2, 6). Consortium MAP#4 improved DSS induced colitis as illustrated by representative gut histology images (D) and ameliorated increases in histopathological score (E&), inflammation (F$) and epithelial injury (G$) compared to vehicle control. Lipocalin-2 concentrations in faeces were increased in mice treated with DSS and concomitant treatment with MAP#4 did not significantly affect lipocalin-2 levels (H&). MAP#4 ameliorated the DSS-induced decrease in goblet cells and intraepithelial lymphocytes, but not the reduced mucin production, when compared to the DSS + vehicle group (I$, J$, K&). MAP#4 was assessed in SKG mouse model of ileitis (S). MAP#4 ameliorated the curdlan-mediated increase in the histopathological score but not the inflammation score when compared to the curdlan + vehicle group (M&, N$). MAP#4 did not significantly improve the curdlan-induced increases in granulomata (O$) and villous distortion score (P$), nor the higher lipocalin-2 levels (Q&). Curdlan treatment resulted in an increase in IL-12p70, IL-6 and IL-23 that was ameliorated by treatment with MAP#4, although this change did not always reach statistical significance when compared to the DSS + vehicle group (R; & for IL-6 and GM-CSF; $ for IL-12). For all data, ns: not significant; *, $p < 0.05$; **, $p < 0.01$; ***, $p < 0.001$; ****, $p < 0.0001$. All data presented as mean and standard deviation. #Two-way ANOVA with Fisher's test for multiple comparison. $Kruskal-Wallis test with uncorrected Dunn's for multiple comparisons. &Uncorrected Brown-Forsythe and Welch ANOVA test with multiple comparisons. %One-way ANOVA with uncorrected Fisher's LSD test for multiple comparison.

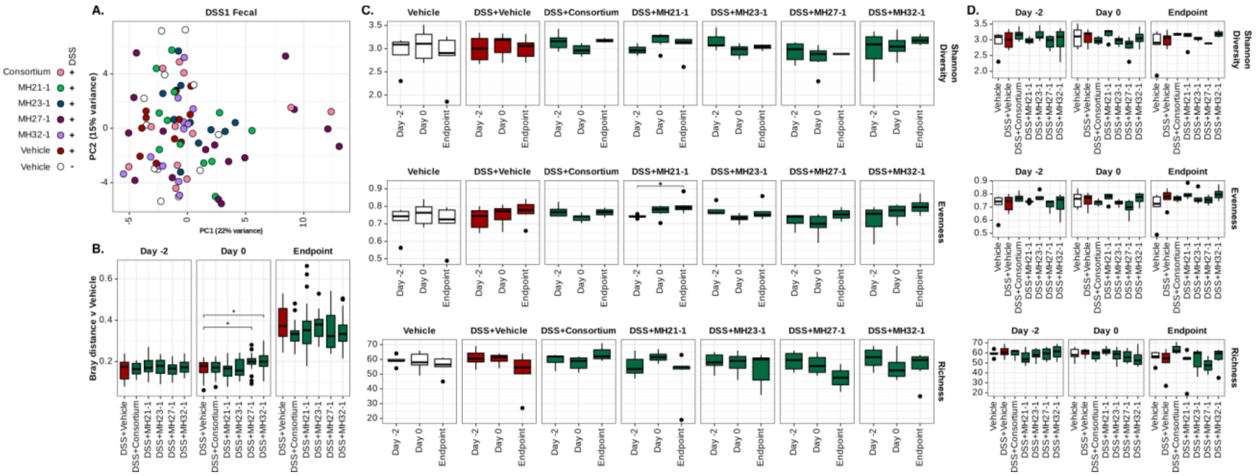

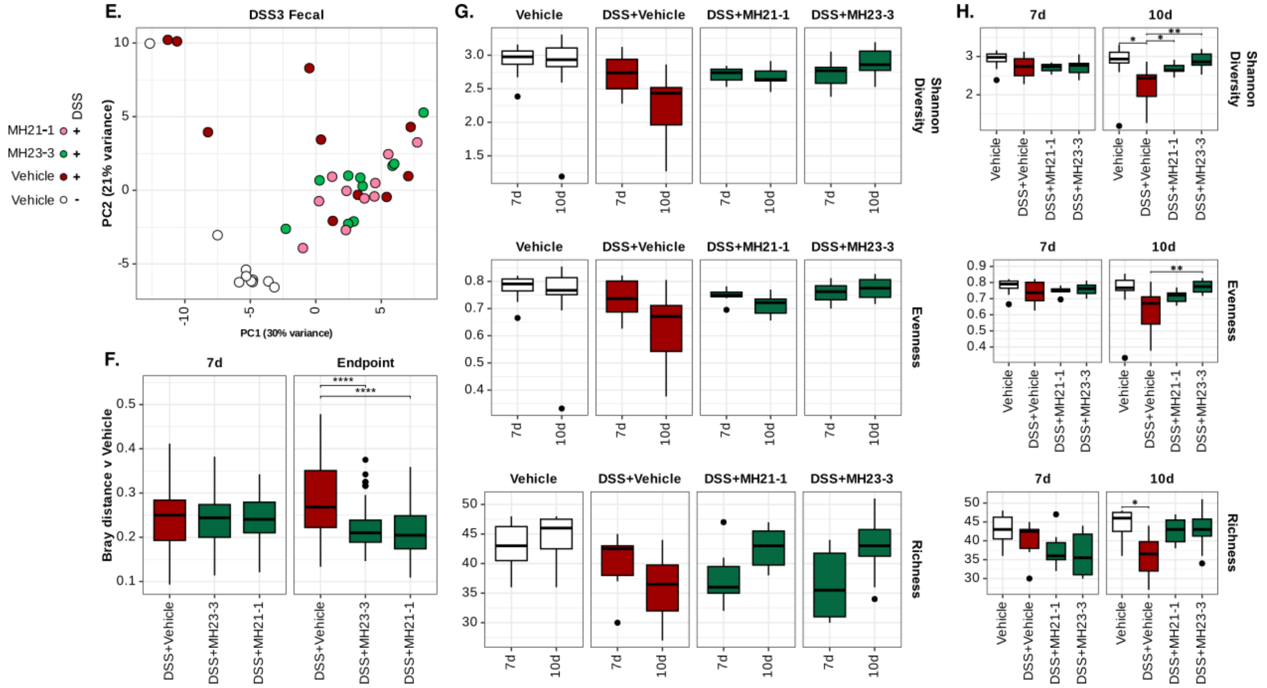


**Supplementary Figure 7**

Metagenome analysis of faecal pellets from the prophylactic (A-D) and therapeutic (E-H) DSS experiments. Significance bars represent FDR corrected P-values of the Wilcoxon Rank Sum Test where * = P-value < 0.05, ** = P-value < 0.01, *** = P-value < 0.001 and **** = P-value < 0.0001. (A) PCA of the Hellinger-transformed species abundance table of faecal pellets collected from mice treated with vehicle and mice treated with DSS and oral gavage of bacterial leads or vehicle. Colours represent each treatment group. (B) Bray-Curtis dissimilarity between mice treated with vehicle and mice treated with DSS and bacterial leads/vehicle. Samples were compared within each timepoint (Day -2, Day 0, and Endpoint). (C) Microbial alpha diversity (Shannon index), evenness and richness of faecal pellet metagenomes, comparing timepoints within each treatment group. (D) As in C, but comparing treatment groups at each timepoint. (E) PCA of the Hellinger-transformed species abundance table of faecal pellets collected from mice treated with vehicle and mice treated with DSS and oral gavage of bacterial leads or vehicle. Colours represent each treatment group. (F) Bray-Curtis dissimilarity between mice treated with vehicle and mice treated with DSS and bacterial leads or vehicle. Samples were compared within each timepoint (Day 7 and Endpoint Day 10). (G) Microbial alpha diversity (Shannon index), evenness and richness of faecal pellet metagenomes, comparing across timepoints within each treatment group. (H) As in G, but comparing treatment groups at each time point.

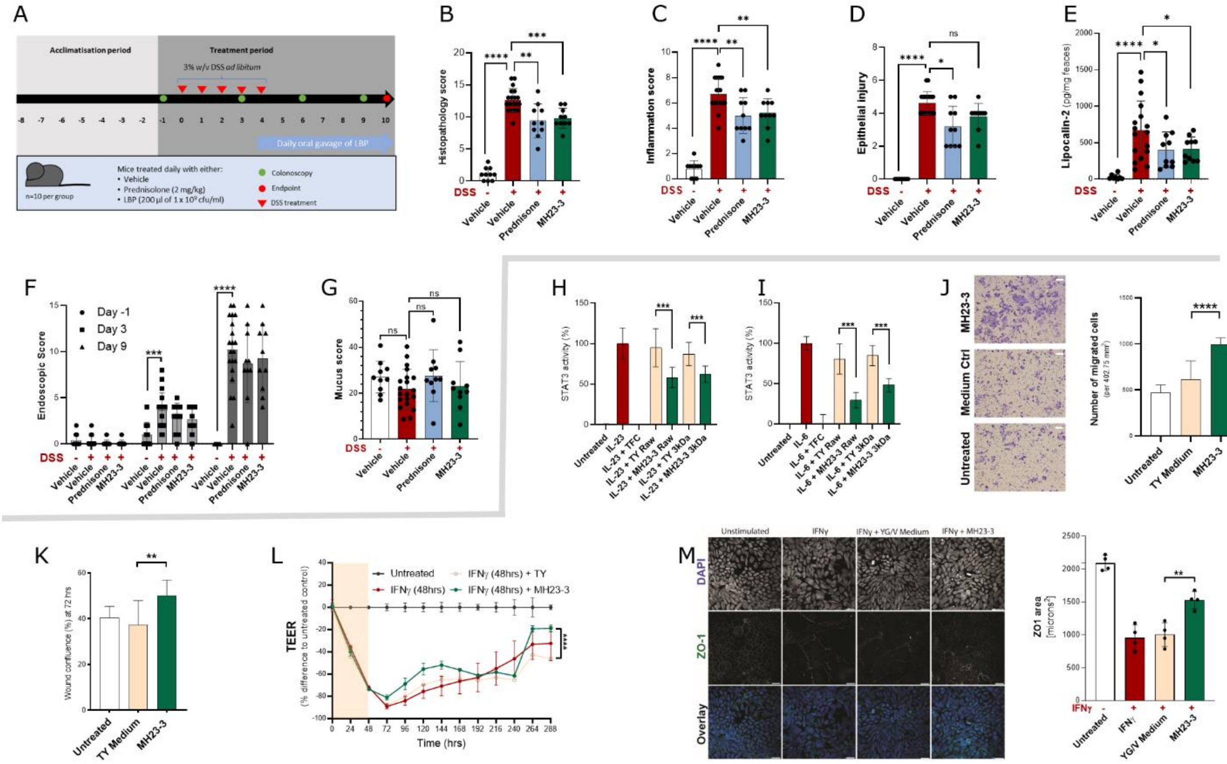


**Supplementary Figure 8.**

Assessment of therapeutic activity of *M. faecis* MH23-3. (A) Study design of the therapeutic DSS mouse model. In this model, MH23-3 ameliorated increases in histopathological score (B[&]), inflammation (C[&]) but not epithelial injury (D[$]) compared to the DSS + vehicle control. MH23-3 ameliorated the increases in faecal lipocalin-2 compared to mice treated with DSS and vehicle (E[&]). MH23-3 did not result in significant improvements in the endoscopic or mucus score compared to mice receiving DSS and vehicle (F[$], G[%]). IL-23-induced STAT3 signalling was inhibited when HEKBlue IL-23 reporter cells were treated with culture supernatant of MH23-3 (H[^]). IL-6-mediated STAT3 signalling was inhibited when HEKBlue IL-6 reporter cells were treated with culture supernatant of MH23-3 (I[^]). MH23-3 extract significantly increased the movement of HCT116 cells compared to the medium extract control in the Transwell migration assay (J[^]; scale bar 100 μm). 72 hours post scratch, HCT116 cells incubated in extract from MH23-3 showed significantly higher wound confluence, compared to cells treated with TY medium extract, in the IncuCyte scratch wound assay (K[^]). T84 gut epithelial cells treated with MH23-3 showed a significantly faster recovery from IFNγ mediated barrier loss compared to the medium control (L[^] at endpoint 288 hrs). MH23-3 ameliorated IFNγ-induced reductions in ZO-1 expression in T84 cells compared to the medium control (M[^]; scale bar 100 μm). For all data, ns: not significant; *, $p < 0.05$; **, $p < 0.01$; ***, $p < 0.001$; ****, $p < 0.0001$. All data presented as mean and standard deviation. [#]Two-way ANOVA with Fisher's test for multiple comparison. [$]Kruskal-Wallis test with uncorrected Dunn's for multiple comparisons. [&]Uncorrected Brown-Forsythe and Welch ANOVA test with multiple comparisons. [%]One-way ANOVA with uncorrected Fisher's LSD test for multiple comparison. [^]T-test of medium control vs lead.

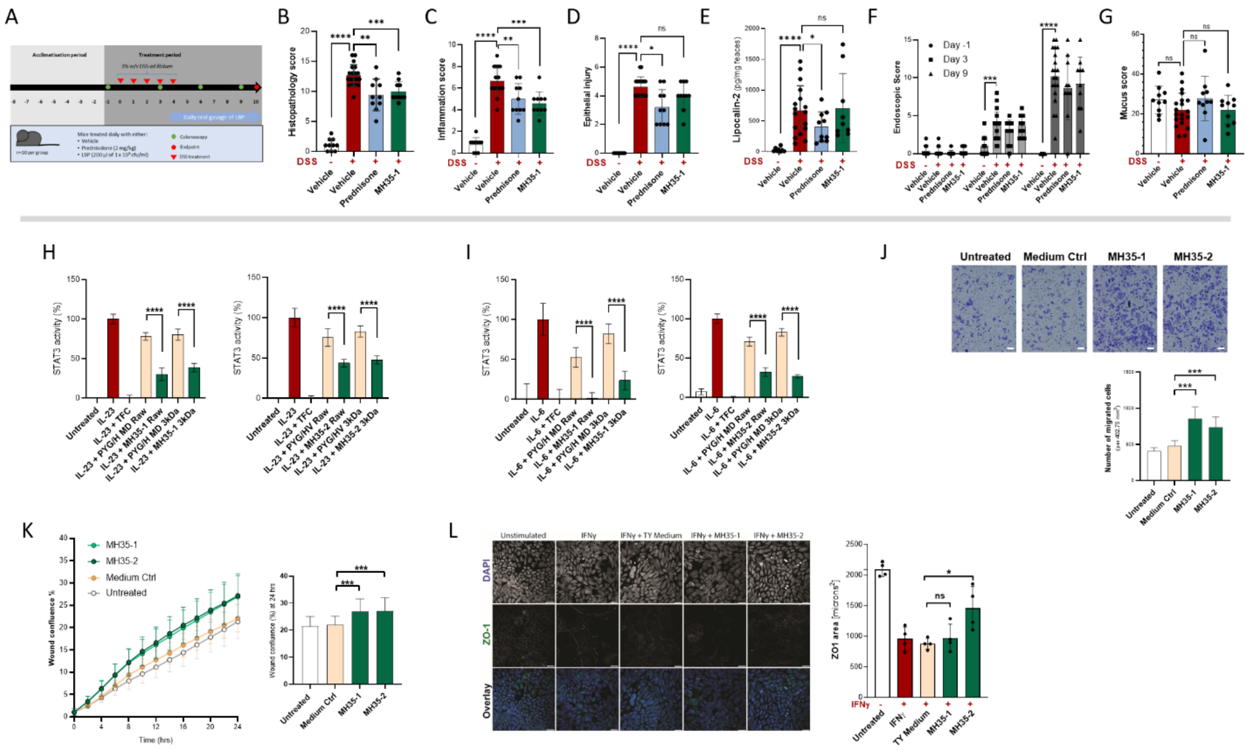


**Supplementary Figure 9.**

Assessment of therapeutic activity of *H. fusiformis* MH35-1 and MH35-2. (A) Study design of therapeutic DSS mouse model. In this model, MH35-1 ameliorated increases in histopathological score (B[&]), inflammation (C[&]) but not epithelial injury (D[$]) compared to the DSS plus vehicle control. Faecal lipocalin-2 levels in mice treated with MH35-1 were comparable to the DSS and vehicle control (E[&]). Daily treatment of DSS-receiving mice with MH35-1 did not result in significant improvements in the endoscopic and mucus score when compared to the DSS + vehicle group (F[$], G[%]). IL-23 mediated STAT3 signalling was inhibited in HEKBlue IL23 reporter cells treated with culture supernatant of MH35-1 and MH35-2 (H[^]). IL-6 induced STAT3 signalling was inhibited in HEKBlue IL6 reporter cells treated with culture supernatant of MH35-1 and MH35-2 (I[^]). MH35-1 and MH35-2 extract significantly increased the movement of HCT116 cells compared to the medium extract control in the Transwell migration assay (J[^]; scale bar 100 μm). 24 hours post scratch, HCT116 cells incubated in extract from MH35-1 and MH35-2 showed significantly higher wound confluence, compared to cells treated with medium extract control, in the IncuCyte scratch wound assay (K[^]). MH35-2, but not MH35-1, ameliorated IFNγ-induced reductions in ZO-1 protein expression in T84 cells compared to the medium control (L[^]; scale bar 100 μm). For all data, ns: not significant; *, $p < 0.05$; **, $p < 0.01$; ***, $p < 0.001$; ****, $p < 0.0001$. All data presented as mean and standard deviation. [#]Two-way ANOVA with Fisher's test for multiple comparison. [$]Kruskal-Wallis test with uncorrected Dunn's for multiple comparisons. [&]Uncorrected Brown-Forsythe and Welch ANOVA test with multiple comparisons. [%]One-way ANOVA with uncorrected Fisher's LSD test for multiple comparison. [^]T-test of medium control vs lead.

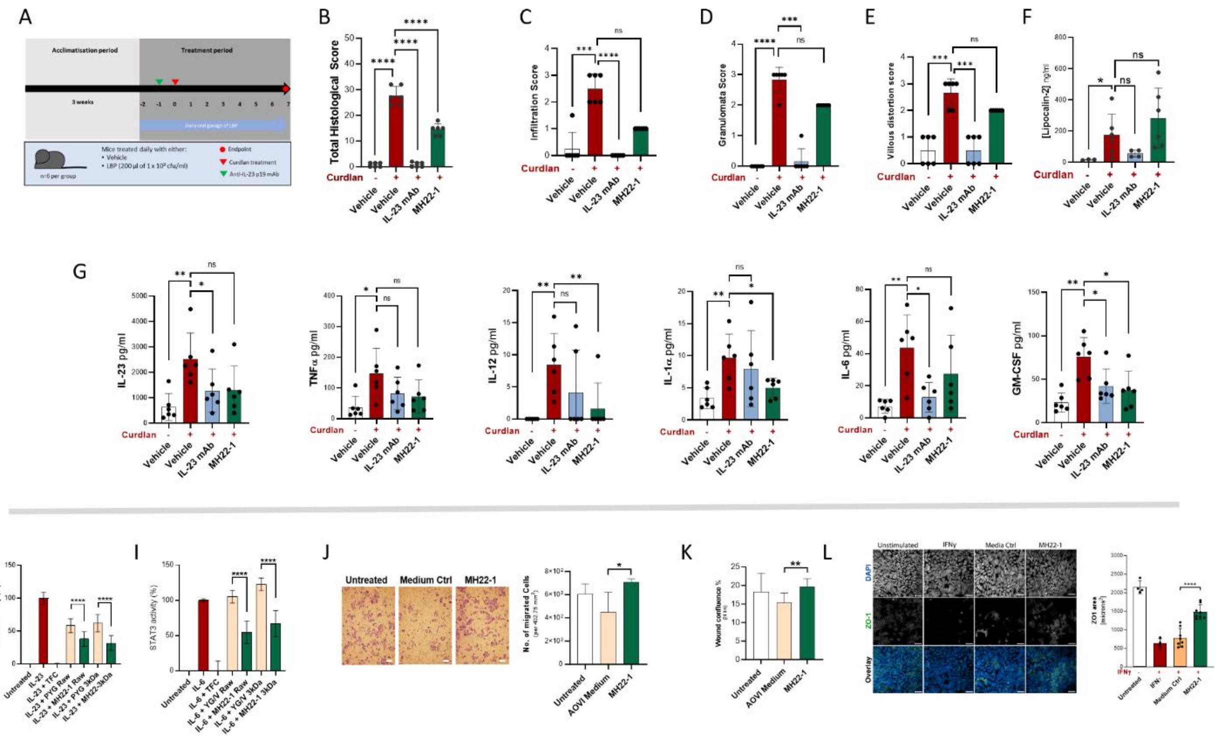


**Supplementary Figure 10.**

Assessment of therapeutic activity of *A. communis* MH22-1. MH22-1 was assessed in the SKG mouse model of ileitis (A). MH22-1 ameliorated the curdlan-induced increase in histopathological score, but not the increase in infiltration score, when compared to the curdlan + vehicle group (B[&], C[$]). MH22-1 did not significantly improve the curdlan-mediated increase in granulomata (D[$]) and villous distortion score (E[$]), nor the higher faecal lipocalin-2 levels (F[&]). Curdlan treatment resulted in a significant increase in cytokines IL-23, TNF, IL-12p70, IL-1α, IL-6 and GM-CSF. Concurrent treatment with MH22-1 caused a significant reduction in IL-12 and GM-CSF levels compared to the curdlan + vehicle group (G[&]) for all cytokines, except IL-12[$]. MH22-1 culture supernatant suppressed IL-23 induced STAT3 signalling in HEKBlue IL-23 reporter cells (H[^]). IL-6-mediated STAT3 signalling was inhibited in HEKBlue IL-6 reporter cells treated with culture supernatant of MH22-1 when compared to the medium control (I[^]). MH22-1 significantly increased the migration of HCT116 cells compared to the medium control in a Transwell migration assay (J[^]; scale bar 100 μm). MH22-1 supernatant extract promoted faster wound closure of HCT116 cells compared to the controls in an IncuCyte scratch wound assay (K[^] 24 hrs post scratch). MH35-2, but not MH35-1, ameliorated IFNγ-induced reductions in ZO-1 protein expression in T84 cells compared to the medium control (L[^]; scale bar 100 μm). For all data, ns: not significant; *, $p < 0.05$; **, $p < 0.01$; ***, $p < 0.001$; ****, $p < 0.0001$. All data presented as mean and standard deviation. [#]Two-way ANOVA with Fisher's test for multiple comparison. [$]Kruskal-Wallis test with uncorrected Dunn's for multiple comparisons. [&]Uncorrected Brown-Forsythe and Welch ANOVA test with multiple comparisons. [%]One-way ANOVA with uncorrected Fisher's LSD test for multiple comparison. [^]T-test of medium control vs lead.

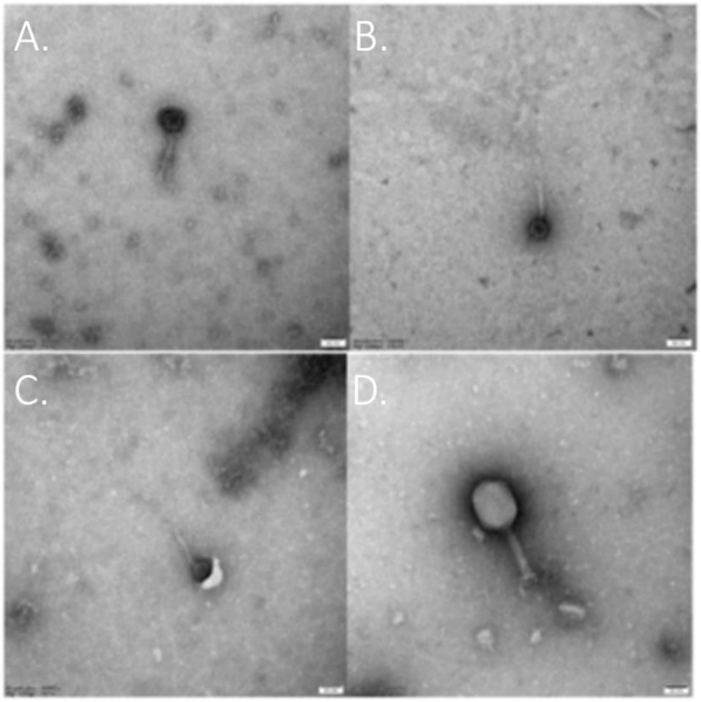


**Supplementary Figure 11.** Phage test results for *G. formicilis* MH32-1. Electron microscopy detected inducible prophage only in MH32-1, which showed high levels of spontaneous induction with lysis of the strain when treated with mitomycin C. A) First round of enrichment of virulent phage test. B) Second round of enrichment of virulent phage test. C) Test for inducible prophage. D) Positive control (T4 phage).

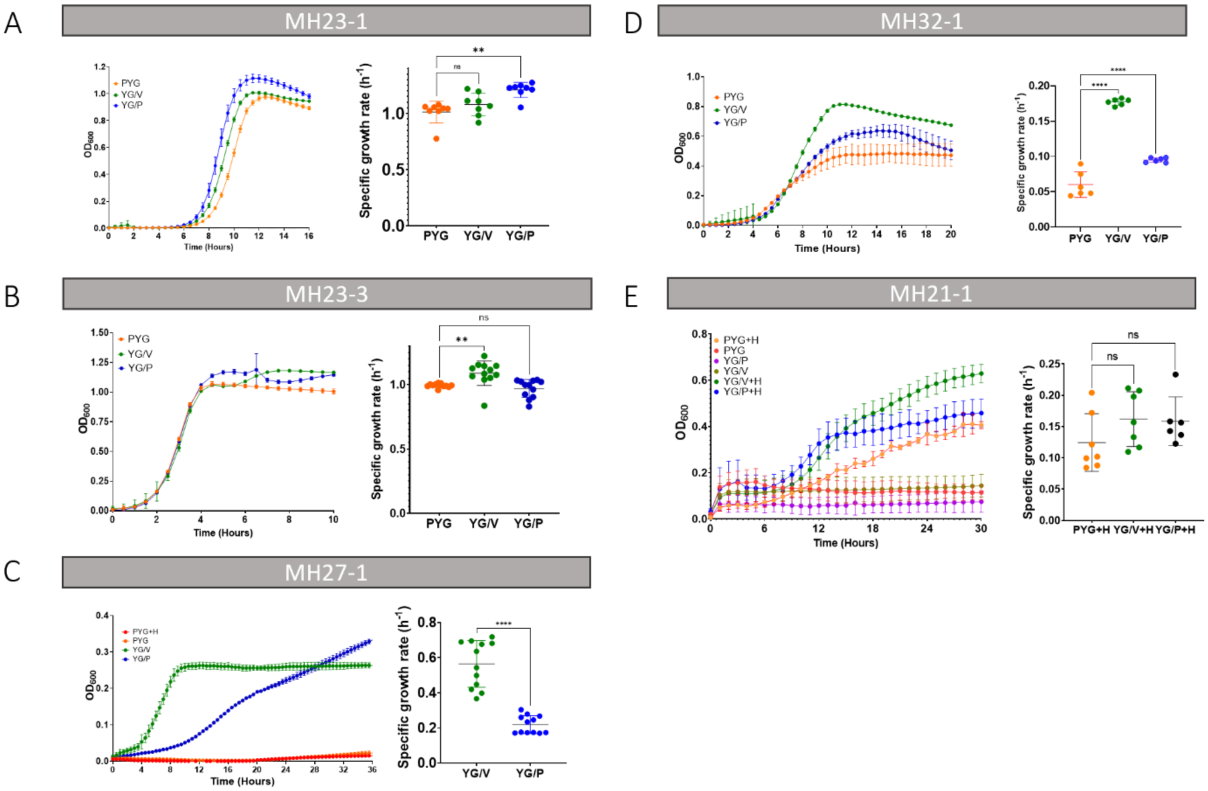


**Supplementary Figure 12.** Growth curve and specific growth rate of *M. faecis* MH23-1 (A), *M. faecis* MH23-3 (B), *H. mulieris* MH27-1 (C), *G. formicilis* MH32-1 (D) and *A. shahii* MH21-1 (E) in animal component containing (PYG) and animal component free (YG/V, YG/P) media. Representative growth curves for single biological replicates are shown. Specific growth rates for all leads except MH32-1 were calculated from 2 biological replicates. *H. mulieris* MH27-1 did not enter exponential growth phase in PYG and so the specific growth rate could not be calculated. *A. shahii* MH21-1 did not enter exponential growth phase in medium lacking haemin and so the specific growth rates could not be calculated.
